# A robust knock-in approach using a minimal promoter and a minicircle

**DOI:** 10.1101/2023.09.15.558008

**Authors:** Margaret Keating, Ryan Hagle, Daniel Osorio-Mendez, Anjelica Rodriguez-Parks, Sarah I Almutawa, Junsu Kang

**Affiliations:** Department of Cell and Regenerative Biology, School of Medicine and Public Health, University of Wisconsin - Madison, Madison, WI, 53705, USA; UW Carbone Cancer Center, School of Medicine and Public Health, University of Wisconsin - Madison, Madison, WI, 53705, USA

**Keywords:** Zebrafish, Knock-in, CRISPR/Cas9, Minicircle, *scn8ab*, *fgf20b*

## Abstract

Knock-in reporter (KI) animals are essential tools in biomedical research to study gene expression impacting diverse biological events. While CRISPR/Cas9-mediated genome editing allows for the successful generation of KI animals, several factors should be considered, such as low expression of the target gene, prevention of bacterial DNA integration, and in-frame editing. To circumvent these challenges, we developed a new strategy that utilizes minicircle technology and introduces a minimal promoter. We demonstrated that minicircles serve as an efficient donor DNA in zebrafish, significantly enhancing KI events compared to plasmids containing bacterial backbones. In an attempt to generate a KI reporter for *scn8ab,* we precisely integrated a fluorescence gene at the start codon. However, the seamlessly integrated reporter was unable to direct expression that recapitulates endogenous *scn8ab* expression. To overcome this obstacle, we introduced the *hsp70* minimal promoter to provide an ectopic transcription initiation site and succeeded in establishing stable KI transgenic reporters for *scn8ab*. This strategy also created a *fgf20b* KI reporter line with a high success rate. Furthermore, our data revealed that an unexpectedly edited genome can inappropriately influence the integrated reporter gene expression, highlighting the importance of selecting a proper KI line. Overall, our approach utilizing a minicircle and an ectopic promoter establishes a robust and efficient strategy for KI generation, expanding our capacity to create KI animals.

## Introduction

Transgenic animals expressing a reporter gene in specific tissues or conditions have critically advanced biomedical research. Traditional transgenic constructs were developed either by subcloning regulatory elements into plasmids or by recombining a reporter gene into bacterial artificial chromosomes (BACs). However, due to the uncertainty of regulatory element location and the limited size capacity of transgenic constructs, DNA sequences required to fully recapitulate endogenous gene expression may not be captured in the transgenic constructs (Chandler et al., 2007; Fuentes et al., 2016). Moreover, transgene fragments randomly integrate into the genome and potentially lead to insertional mutagenesis (Espinoza et al., 2019; Goodwin et al., 2019). Therefore, targeted integration is optimal to create knock-in (KI) reporter animals.

CRISPR/Cas9-mediated genome editing is a revolutionized technique that enables the insertion of DNA of interest at a targeted genomic locus (Cong et al., 2013; Hwang et al., 2013; Mali et al., 2013; Wang et al., 2013; Yang et al., 2013). A donor DNA template (exogenous DNA cassette) can be integrated into the genome when a double-strand break (DSB) induced by CRISPR/Cas9 is repaired. Donor DNAs are typically plasmids containing bacterial backbone sequences, 5’ and 3’ homology arms (HAs), and sgRNA site(s) (Ata et al., 2016; Hoshijima et al., 2016; Li et al., 2016). To successfully generate a KI line, several factors should be considered, such as in-frame editing, amplifying the faint expression of the target gene to visualize the signal, and preventing the integration of bacterial DNA, which can hinder transcription in eukaryotic cells (Ata et al., 2018; Riu et al., 2005; Wierson et al., 2020). To circumvent these limitations, we established a new strategy by utilizing minicircle technology and introducing an ectopic promoter. Minicircle DNA vectors, in which the bacterial backbone is robustly removed *in vitro* (Kay et al., 2010), can greatly increase KI efficiency. Moreover, we demonstrated that incorporating an ectopic minimal promoter into a donor vector provides a remarkable benefit to direct reporter expression of the target genes. In situations when targeting the endogenous promoter to trigger reporter expression proves ineffective, this approach will be especially advantageous. Consequently, our effective strategies expand our capacity to create KI animals.

## Material and Methods

### Key resources table

| REAGENT or RESOURCE | SOURCE | IDENTIFIER |
| --- | --- | --- |
| Chemicals, peptides, and recombinant proteins |  |  |
| Tricaine methanesulfonate | Pentair | TRS1 |
| Experimental models: Organisms/strains |  |  |
| Zebrafish: Ekkwill: Wild type | Ekkwill Breeders |  |
| Zebrafish: AB: Wild type | AB, Breeders |  |
| Zebrafish: *scn8ab*<sup>temca or pd291</sup> | 17 |  |
| Zebrafish: *scn8ab*<sup>EGFP</sup> | This paper |  |
| Zebrafish: *scn8ab*<sup>hsp70pro:EGFP</sup> Line 2 (*uwk19*) and 13 (*uwk20*) | This paper |  |
| Zebrafish: *fgf20b*<sup>hsp70pro:EGFP</sup> Line 1 (*uwk21*) and 3 (*uwk22*) | This paper |  |
| *E.coli*: ZYCY10P3S2T | SBI | MN900A-1 |
| Oligonucleotides |  |  |
| See **Table S2** for subcloning primers | This paper | N/A |
| Recombinant DNA |  |  |
| Plasmid: pMC.BESPX-MCS2 | SBI | MN100B-1 |
| Plasmid: pMC.BESPX-EGFP-pA | This paper | N/A |
| Plasmid: pMC-EGFP-SV40pA-FRT-acry:mcherry-FRT | This paper | N/A |
| Plasmid: pMC-*scn8ab*:EGFP-SV40pA-FRT-acry:mcherry-FRT | This paper | N/A |
| Plasmid: pMC-*hsp70pro:scn8ab*-EGFP-SV40pA-FRT-acry:mCherry-FRT | This paper | N/A |
| Plasmid: pMC-*hsp70pro:scn8ab*-EGFP-SV40pA-FRT-acry:mCherry-FRT two sgRNA sites | This paper | N/A |
| Plasmid: pMC-*fgf20b*:EGFP-SV40pA | This paper | N/A |
| Plasmid: pMC-*hsp70pro:fgf20b*:EGFP-SV40pA | This paper | N/A |
| Plasmid: pMC-*hsp70pro:fgf20b*:EGFP-SV40pA two sgRNA sites | This paper | N/A |

| Software and algorithms |  |
| --- | --- |
| Zeiss ZEN | Carl Zeiss |
| NIS-Elements | Nikon |
| Image J/FIJI |  |
| Photoshop | Adobe |

### Zebrafish maintenance and procedures

Wild-type or transgenic male and female zebrafish of the outbred Ekkwill (EK) or AB strains ranging up to 18 months of age were used for all zebrafish experiments. The water temperature for adult animals was maintained at 26°C unless otherwise indicated. Embryos and larvae were maintained at 28°C in egg water containing 300 mg/L sea salt, 75 mg/L calcium sulfate, 37.5 mg/L sodium bicarbonate, and 0.0001% methylene blue. Animals were anesthetized in 0.02% tricaine until gill movement stopped. To assess locomotion defect, compound heterozygotes of *scn8ab^hsp70:EGFP^* and *temca* and control siblings were placed at 33°C hot water and their swimming behavior was recorded using a smartphone camera. Work with zebrafish was performed in accordance with University of Wisconsin-Madison guidelines.

### Subcloning and minicircle generation

pMC.BESPX-*EGFP-pA* was generated by inserting an *EGFP-pA* cassette into the pMC.BESPX-MCS2 (MN100B-1, SBI) via NheI/XbaI and BstBI enzymes-mediated subcloning. pMC-*EGFP-SV40pA-FRT-acry-mCherry-FRT* was generated by inserting *FRT-a-cry:mCherry-FRT* cassette via ClaI and BamHI enzymes-mediated subcloning into pMC.BESPX-*EGFP-pA* vector. 5’ and 3’ HAs were amplified using genomic DNA extracted from fish that were used for injection. The corresponding sgRNA site for *scn8ab* and *fgf20b* was placed at the 5’ end of 5’ HA. sgRNA sequences for *scn8ab* and *fgf20b* are “GGGCC TGGAG GTGCG AGCAG” and “GGAAT AAATG TAGGT TAGGT”, respectively. These HA fragments were inserted into pMC-*EGFP-SV40pA-FRT-acry:mcherry-FRT* and pMC.BESPX-*EGFP-pA* for *scn8ab* and *fgf20b*, respectively. *hsp70* promoter fragment was added via Gibson assembly or enzyme-based subcloning for *scn8ab* or *fgf20b*, respectively. To purify the minicircle vectors, parental plasmids were transformed into the ZYCY10P3S2T *E. coli* Minicircle producer strain (MN900A-1, SBI). Minicircles were prepared as described in the **Supplementary Information**. To prepare plasmids used in **Fig. 2D** and **4E**, we added additional sgRNA site at the 3’ end of 3’ HA, resulting in two sgRNA sites in the plasmids. Inside of cells after injection, these donor plasmids provided a similar size of donor template as a result of CRISPR/Cas9-mediated cut, compared to minicircle. Primers used for subcloning are listed in **Supplementary Table S2**.

### Generation of *scn8ab* and *fgf20b* KI reporter lines and RT-PCR

To generate KI reporter lines, CRISPR sgRNA target sites were designed using the CRISPRscan web tool (https://www.crisprscan.org/) (Moreno-Mateos et al., 2015). sgRNAs were synthesized by a cloning-free method as described in (Varshney et al., 2015). Briefly, primers containing T7 promoter, sgRNA target and partial sgRNA scaffold sequences were annealed with the universal sgRNA 3’ scaffold primer, and the annealed oligos were filled using the thermocycler and PCRBio polymerase (Genesee Scientific) to generate sgRNA templates. sgRNAs were synthesized using the HiScribe T7 kit (NEB, E2050S) and purified by the RNA purification Kit (Zymogen, R1016) according to the manufacturer’s instructions. A sgRNA (25∼30 ng/ul) and a donor minicircle or a plasmid (20-25 ng/ul) were mixed and co-injected with Cas9 protein (0.5ug/ul; PNABio, CP01) into the one-cell stage embryos. Genomic DNA was extracted, and KI was confirmed by genotyping. Primers used for genotyping are listed in **Table S2**. EGFP positive larvae were sorted and raised to adulthood, and founders were screened with F_1_ progenies. The founder was outcrossed with wild-type animals to generate heterozygous animals. RNA was isolated from brain tissues of wild-type and *scn8ab^EGFP^* fish using Tri-Reagent (ThermoFisher). Complementary DNA (cDNA) was synthesized from 300 ng to 1 μg of total RNA using a NEB ProtoScript II first strand cDNA synthesis kit (NEB, E6560). The sequences of the primers used for RT-PCR are listed in **Table S2**.

### Imaging, immunostaining, and histology

Whole-mount larval images were acquired using an AxioZoom stereo fluorescence microscope (Zeiss) or a Nikon A1R-s confocal laser scanning microscope (Nikon). For confocal images of the adult brain, maximum projections were obtained using approximately 100 slices every 2 µm apart. Further image processing was carried out manually using Zen (Zeiss), NIS-Elements (Nikon), Photoshop, or FIJI/ImageJ software. For making frozen sections, brains were fixed with 4% PFA overnight at 4°C, then washed with PBST (0.1% Tween-20 in PBS) 3 times for 10 minutes. Tissues were then embedded in 1.5% agarose with 5% sucrose and incubated in 30% sucrose overnight at 4°C. Frozen blocks were sectioned at 16 µm in a cryostat. The primary and secondary antibodies used in this study were as follows: anti-HuC/D (mouse, clone 16A11, #A21271, Invitrogen, 1:100), anti-GFP (chicken, AB_2307313, Aves Labs, 1:2000), Alexa Fluor 488 (goat anti-chicken, #A11039, Life Technologies, 1:500) and Alexa Fluor 594 (goat anti-mouse, #A11005 Life Technologies, 1:500). *in situ* hybridization of 4% paraformaldehyde-fixed larvae was performed as previously described (Schiavo and Tamplin, 2022). To generate digoxigenin-labeled probes for *scn8ab*, we used 3’ untranslated region of *scn8ab* cDNA, which does not show conservation with other *scn* genes(Novak et al., 2006). To generate digoxigenin-labeled probes for *fgf20b*, we used full length of *fgf20b* cDNA. Primer sequences are listed in **Table S2**.

### 5’ RACE

We used Rapid amplification of cDNA ends (RACE) to identify the 5’ end of the EGFP transcript (5’ RACE). The first step involved generating cDNAs using an EGFP reverse transcription (RT) primer and a universal sequence introduced to the 3’ end of the cDNA by a template switching oligo (TSO). We designed an RT primer 114bp downstream of the EGFP start site (TTCAGGGTCAGCTTGCCGTAGGT). After extracting RNA from *scn8ab^hsp70:EGFP^* line 2 and *fgf20b^hsp70:EGFP^* line 3, 600ng of RNA, the EGFP RT primer (1µM final), and dNTP (1mM final) were annealed using 70°C for 5 minutes. A mixture of template switching buffer, template switching oligo (3.75µM final, (GCTAATCATTGCAAGCAGTGGTATCAACGCAGAGTACATrGrGrG), and RT enzyme mix (NEB, M0466) was combined with the annealed mixture. The reaction was then incubated at 42°C for 90 minutes, and heat inactivated at 85°C for 5 minutes.

Next, the 5’ end of the transcript was amplified via PCR. We designed an EGFP specific PCR primer 65bp downstream of the EGFP start site (CCGGACACGCTGAACTTGTGGCCGTTTACG) and a primer specific to the TSO handle (CATTGCAAGCAGTGGTATCAAC). The generated cDNA was diluted 2x and 1µL was used for PCR amplification using PCR Biosystems VeriFi Mix (Genesee Scientific, 17-308B). A touchdown-PCR method was used for amplification. We used 5 cycles of a 72°C annealing temperature followed by 5 cycles of a 70°C annealing temperature. The PCR then ran as normal for 30 cycles using a 65°C annealing temperature. The PCR product was extracted via gel electrophoresis and purified. We found two bands for each line and sent both for Sanger sequencing using an EGFP primer 45bp downstream of the EGFP start site (CGTCGCCGTCCAGCTCGACCAG).

## Results

### Targeting endogenous *scn8ab* promoter fails to direct reporter gene expression

Our previous forward genetic screening identified the *temca* mutant which exhibits temperature-sensitive locomotion and fin regeneration defects (Osorio-Mendez et al., 2022). The mutated gene was defined as *scn8ab* that encodes a voltage-gated sodium channel, Nav1.6. Although previous studies examined *scn8ab* expression by in situ hybridization (ISH) (Novak et al., 2006; Osorio-Mendez et al., 2022), its spatiotemporal expression pattern was unexplored by transgenic assay. To address this, we attempted to generate a *scn8ab* KI reporter line using the CRISPR/Cas9 technique. We designed a donor vector to integrate the EGFP reporter gene at the translation initiation site (TIS) of *scn8ab,* located in exon 2 (**Fig. 1A**). The donor plasmid consists of 5’ and 3’ homology arms (HAs), sgRNA site, EGFP, and SV40 polyA signal (pA) (**Fig. 1A**). sgRNA was selected from the genomic DNA between 5’ and 3’ HAs, preventing continuous cleavage that may cause unexpected editing. The 5’ HA is a 518-bp fragment upstream of the *scn8ab* TIS and the selected *scn8ab* sgRNA site was added at the 5’ end of the 5’ HA, which is used for linearization in a cell after injection. The 534-bp 3’ HA fragment starts after the sgRNA site. As *scn8ab temca* mutants display phenotypes during juvenile but not larval stages (Osorio-Mendez et al., 2022) and its transcript expression in larvae is highly restricted to a few cells of trigeminal ganglia and RB cells (Novak et al., 2006), we expected undetectable or limited *scn8ab* expression in larvae. Thus, we added an FRT-flanked lens mCherry selection marker, which facilitated the selection of larvae carrying the modified genome. The final donor DNA was prepared via minicircle technology (Kay et al., 2010), which produces a minicircle containing necessary DNAs without the bacterial backbone sequences. Consequently, a single sgRNA site in the minicircle is sufficient to provide a linearized, bacterial DNA-absent cassette inside of the cell, compared to two sgRNA sites for the bacterial plasmid (**Fig. S1**).

**Figure 1.**
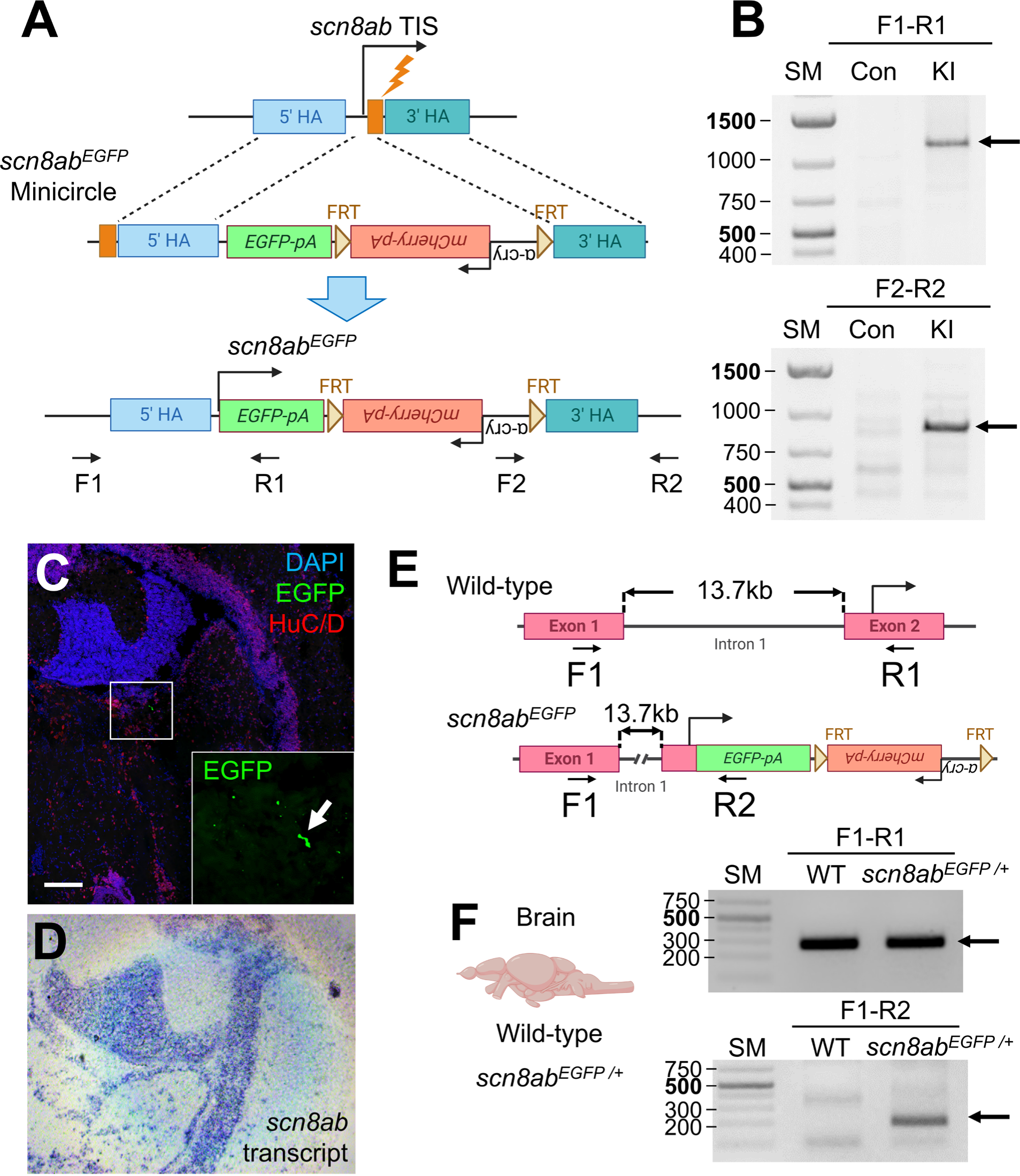
Precise integration of *EGFP* at *scn8ab* start codon fails to direct gene expression of endogenous *scn8ab*. (**A**) Schematic of the strategy to integrate *EGFP* at the *scn8ab* start codon using a minicircle donor. TIS, translation initiation site (**B**) PCR analysis to genotype upstream and downstream regions of the integrated site at the *scn8ab* locus. Corresponding primers are distally located to the HA as shown in (**A**). Arrows indicate correctly amplified fragment. SM, size marker. (**C**) Immunofluorescence EGFP signal of *scn8ab^EGFP^* in brain tissues. Scale bar, 100µm. (**D**) *in situ* hybridization (ISH) signal of *scn8ab* in brain tissues. (**E**) Schematic of the zebrafish *scn8ab* locus and inserted EGFP. (**F**) RT-PCR analysis of *scn8ab* and *EGFP* transcript expression in brain of wild-type (WT) and *scn8ab^EGFP^* heterozygotes. Corresponding primers are shown in (**E**).

After co-injection of the donor minicircle, *scn8ab* sgRNA, and Cas9 protein into the one-cell-stage embryos, we observed lens mCherry expression at 5 days post-fertilization (dpf), but EGFP was undetectable. Based on lens mCherry expression, we established one stable line, of which seamless integration was confirmed by PCR followed by Sanger sequencing (**Fig. 1B and Fig. S2, 3**). Despite a seamless integration of the reporter gene, EGFP expression was undetectable in larvae and extremely limited in adults. Immunostaining was conducted against EGFP in the adult brain to amplify a weak signal, but a few neurons were labeled (**Fig. 1C**). Although our ISH analysis showed broad expression of *scn8ab* transcript in the brain (**Fig. 1D**), EGFP immunofluorescence was detectable only in a few cells of *scn8ab* KI brain (**Fig. 1C**). The first exon of the *scn8ab* isoform is located ∼13.7 kb upstream from the second exon, where the TIS is placed (**Fig. 1E and Fig. S2**). We evaluated the presence of the *EGFP* transcript in the cDNA libraries generated with adult brain tissues using RT-PCR followed by Sanger sequencing. Our analysis confirmed that brain tissues can generate a transcript containing EGFP at the *scn8ab* locus (**Fig. 1F**), indicating that the integrated *EGFP* cassette is transcribed, although it appears to be highly restricted to a few cells, or may not be translated due to unknown reasons.

### A minimal promoter and a minicircle donor facilitate the generation of *scn8ab* knock-in reporter line

The *heat shock 70* (*hsp70*) minimal promoter, hereafter referred to as *hsp70pro,* was previously used to generate enhancer trap lines in which enhancers in close proximity to the integration site can interact with *hsp70pro* to direct downstream cassette expression, such as a fluorescence protein (Asakawa et al., 2008; Nagayoshi et al., 2008). *hsp70pro* was also employed to create KI reporter lines as it can enhance the efficiency of reporter expression (Hawkins et al., 2021; Kimura et al., 2014). To determine whether *hsp70pro* can drive *EGFP* expression when integrated at the *scn8ab* genomic locus, we added *hsp70pro* upstream of the *EGFP* start codon (**Fig. 2A, B**). Remarkably, the *hsp70pro*-containing construct directed expression in the neurons of larvae, implying that *hsp70pro* can circumvent the problem of the original construct lacking an ectopic promoter (**Fig. 2B, C**). In addition to neuronal expression, we observed weak to modest EGFP expression in some skeletal muscle cells, which may be driven by *hsp70pro* as previously reported (Asakawa et al., 2008; Kimura et al., 2014; Nagayoshi et al., 2008). (**Fig. 2B**). Next, we compared the KI efficiency of the donor DNA between a plasmid and a minicircle. A plasmid contains two sgRNA sites flanking 5’ and 3’ HA, resulting in a similar size of donor template inside of cells after injection, compared to minicircle. While a plasmid donor yielded 28.4% efficiency (19/67 larvae), a minicircle donor exhibited a 110% higher efficiency (59.8%; 259/433 larvae) (**Fig 2D**). Thus, our results demonstrate that a minicircle and *hsp70pro* are beneficial additions to facilitate the generation of a KI reporter line.

**Figure 2.**
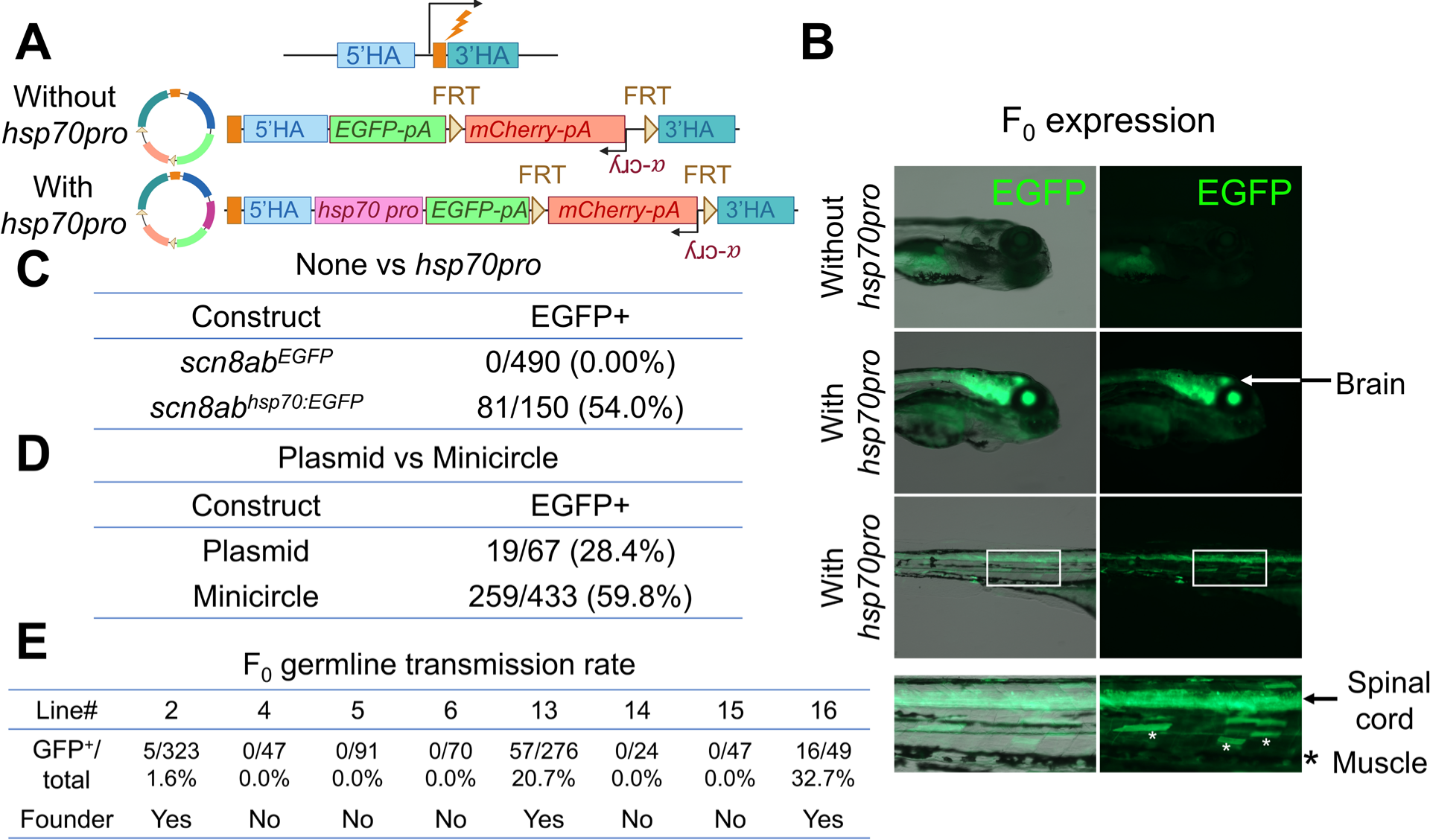
Generation of *scn8ab^hsp70:EGFP^* reporter line using a minicircle donor and an ectopic *hsp70* promoter (*hsp70pro*). (**A**) Schematic of the strategy to integrate *EGFP* or *hsp70:EGFP* at the *scn8ab* start codon using minicircle donor. (**B**) Representative images of F_0_ larvae injected with *EGFP* (without *hsp70pro*) or *hsp70pro:EGFP* (with *hsp70pro*) minicircle donor. (**C**) Comparison of EGFP expressing larvae after injecting *EGFP* vs *hsp70pro:EGFP* minicircle donor. (**D**) Comparison of EGFP expressing larvae after injecting *hsp70pro:EGFP* plasmid or minicircle donor. (**E**) Screening results and germline transmission rate to establish the *scn8ab^hsp70:EGFP^* lines.

### *scn8ab^hsp70:EGFP^* drives neuronal expression that recapitulates the endogenous *scn8ab* gene

To determine whether our KI approach can successfully generate a stable line, EGFP^+^ embryos were raised to adulthood and mated with wild-type to examine their progenies. Among the 8 F_0_ screened fish, we found three stable lines displaying neuronal EGFP induction in larvae with low to modest germline transmission rates (1.6%, 26.0%, and 32.7% for Line 2, 13, and 16, respectively) (**Fig. 2E**). All three of the established *scn8ab^hsp70:EGFP^* lines direct neuronal expression (**Fig. 3A**) with variable skeletal muscle expression patterns and intensities. To characterize the feature of non-specific muscle expression, we sorted heterozygote Line 2 larvae that displayed EGFP expression exclusively in neurons, raised them to adulthood, outcrossed them with wild-type, and analyzed the expression pattern in progenies. We found that neuronal expression is consistent and strong through all EGFP^+^ larvae. We did not observe any larvae exhibiting EGFP expression exclusively in muscle tissue. As skeletal muscle displayed variable expression results, we could score muscle expression as “none”, “weak”, “moderate”, and “strong” and were unable to find a consistent ratio among multiple mating pairs (**Fig. 3B and Table S1**). Notably, most *scn8ab^hsp70:EGFP^* Line 2 larvae were scored as weak, having highly restricted EGFP muscle expression to a few cells throughout the body. This weak expression in skeletal muscle became visible when brightness and intensity were increased, otherwise it was not observable. We sorted “none” or “weak” larvae and assessed EGFP expression in adults. Whole-mount imaging of the trunk in adults displayed obvious neuronal EGFP expression, but muscular EGFP expression was undetectable (**Fig. 3C**). EGFP also widely labels neurites in adult brain tissues, mimicking endogenous *scn8ab* expression (**Fig. 3D**). Thus, we established *scn8ab^hsp70:EGFP^* lines that faithfully capture the endogenous expression of *snc8ab*. Our data show that leaky induction of *hsp70pro* drives variable reporter gene expression in skeletal muscle due to stochastic transcription during embryogenesis, but careful selection of fish that exhibit minimal non-specific expression can mitigate the influence of this unfavorable effect.

**Figure 3.**
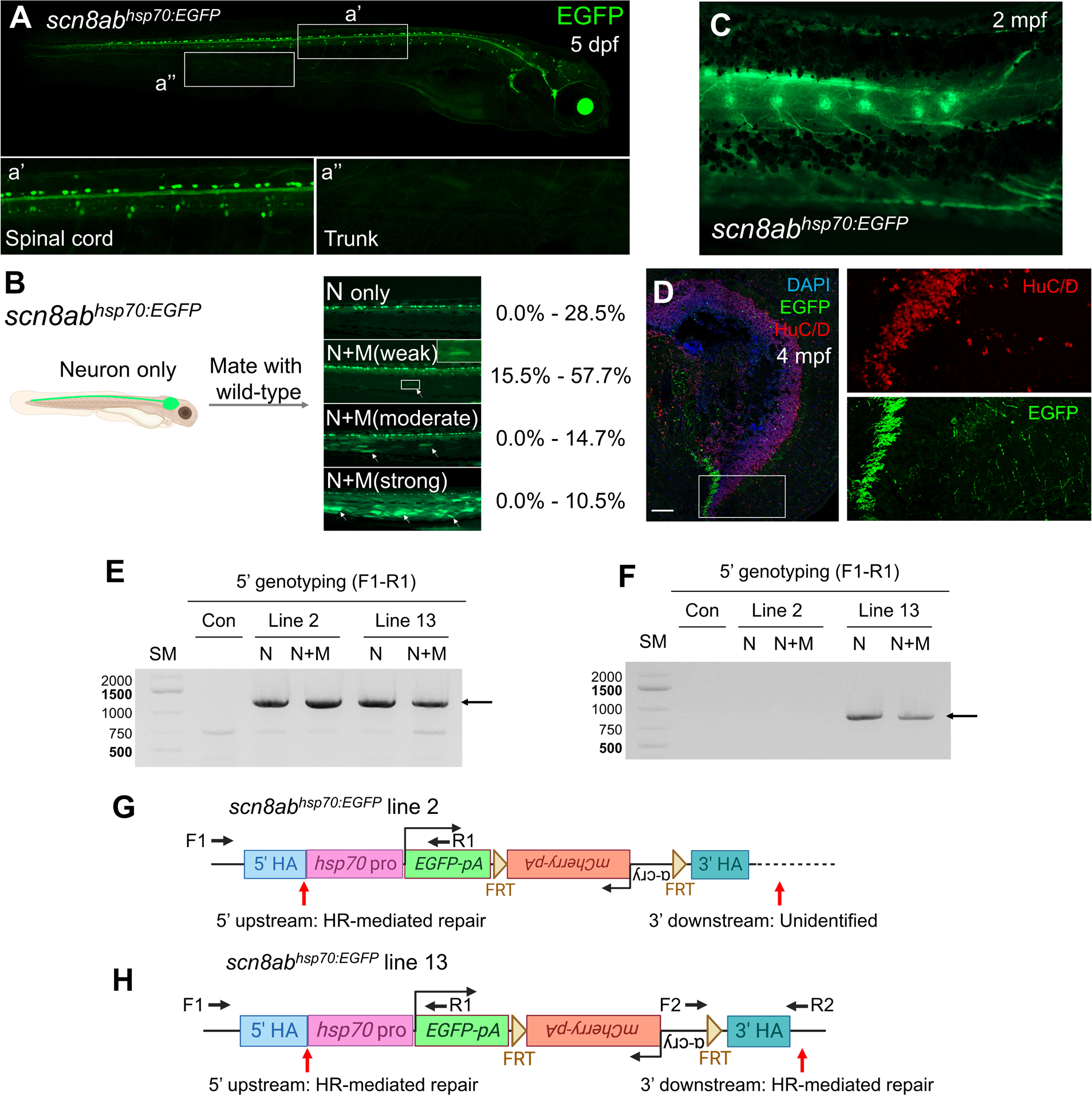
Characterization of *scn8ab^hsp70:EGFP^*. (**A**) Representative image of the *scn8ab^hsp70:EGFP^* stable Line 2 at 5 dpf. (**B**) Variability of skeletal muscle expression of *scn8ab^hsp70:EGFP^* Line 2. Representative images of variable EGFP expression pattern for *scn8ab^hsp70:EGFP^* and a range of the ratio are indicate in the right. N only, neurons without skeletal muscle expression. N+M (weak), neurons plus limited EGFP expression in a few skeletal muscle cells. N+M (moderate), neurons plus modest EGFP expression in some skeletal muscle cells. N+M (strong), neurons plus strong EGFP induction in a substantial number of skeletal muscle cells. (**C**) Representative whole-mount image for the body trunk of *scn8ab^hsp70:EGFP^* stable Line 2 at 2 month post-fertilization (mpf). Neuronal EGFP in the spinal cord and dorsal root ganglia are visible. Muscular EGFP expression is likely undetectable. (**D**) Immunofluorescence EGFP signal of *scn8ab^hsp70:EGFP^* in brain tissues obtained from 4 mpf. Scale bar, 100µm. HuC/D labels neuronal nuclei. (**E, F**) PCR analysis to genotype upstream (**E**) and downstream (**F**) regions of the integrated site at the *scn8ab* locus. Corresponding primers are distally located to the HA as shown in **(G, H)**. Arrows indicate correctly amplified fragment. (**G, H**) Schematic of edited genomic DNA sequence for *scn8ab^hsp70:EGFP^* line 2 (**G**) and 13 (**H**).

We explored how the *scn8ab* genome is edited in two of *scn8ab^EGFP^* KI lines. Genomic DNA was collected from the neuron (Neu) and neuron + muscle (Neu+Mus) groups of Line 2 and Line 13. 5’ upstream and 3’ downstream regions of the target site were amplified with corresponding primer pairs. PCR analyses followed by Sanger sequencing revealed that the 5’ upstream region is integrated via homologous recombination (HR)-mediated repair for both lines (**Fig. 3E-H and S4**). For the 3’ downstream region, PCR failed to amplify any specific DNA bands for Line 2 (**Fig. 3F, G**), indicating a potential large deletion. In contrast, PCR analysis with Line 13 generated the expected size band, the sequencing of which indicates HR-mediated repair (**Fig. 3F, H and S4**). Since edited genomic DNA disrupts the *scn8ab* coding region, the *scn8ab^EGFP^* allele likely results in a loss-of-function of *scn8ab*. Indeed, our genetic validation demonstrated compound heterozygotes of Line 2 with *temca* exhibit locomotion defect at 33°C, indicating that the EGFP cassette is correctly integrated at the *scn8ab* locus (**Video S1**).

### Generation of *fgf20b* KI reporter line

*fgf20b* is known to play a role in the development of the head skeletal system, including pharyngeal cartilage (Yamauchi et al., 2011). Consistent with this, our ISH can detect *fgf20b* transcript in the pharyngeal arches at 1 dpf (**Fig. 4B**). To further investigate the spatiotemporal expression domain of *fgf20b*, we applied our strategy to create a *fgf20b* KI reporter line. First, we prepared a donor minicircle containing 632-bp 5’ HA containing the sgRNA site, *EGFP* reporter gene, SV40-pA, and 436-bp 3’ HA. Unlike the *scn8ab* design, the sgRNA site for *fgf20b* was located at the 5’ end of 5’ HA in the genome (**Fig. S5**). Previous studies demonstrated that a sgRNA located in the 5’ HA also can be employed to establish KI reporter lines (Kesavan et al., 2017; Kimura et al., 2014; Li et al., 2015). None of the larvae co-injected with the *fgf20b* donor minicircle, *fgf20b* sgRNA, and Cas9 showed EGFP expression (**Fig. 4C, D**). To circumvent this problem, we added the *hsp70pro* upstream of *EGFP*, which directed strong EGFP expression in the pharyngeal arch region (33.94%; 92/271 larvae) (**Fig. 4C, D**). Similar to the *scn8ab* KI reporter line, we observed the *hsp70pro*-containing construct directed EGFP expression in a few skeletal muscle cells (**Fig. 4C**) which was not observed by ISH analysis (**Fig. 4B**). We also demonstrated that the KI efficiency of the minicircle is higher than that of the plasmid as a donor template by 267% enhancement (4.5% and 16.5% for plasmid and minicircle, respectively) (**Fig. 4E**). Among 4 screened F_0_ fish, 2 fish were identified to transmit the EGFP to the next generation with around 25% germ line transmission rate (**Fig. 4F and 5A**). EGFP is also detectable in unidentified cells of the head at the adult stage (**Fig. S5**). Therefore, we demonstrated that our strategy significantly improved the potency and efficiency of KI reporter generation.

**Figure 4.**
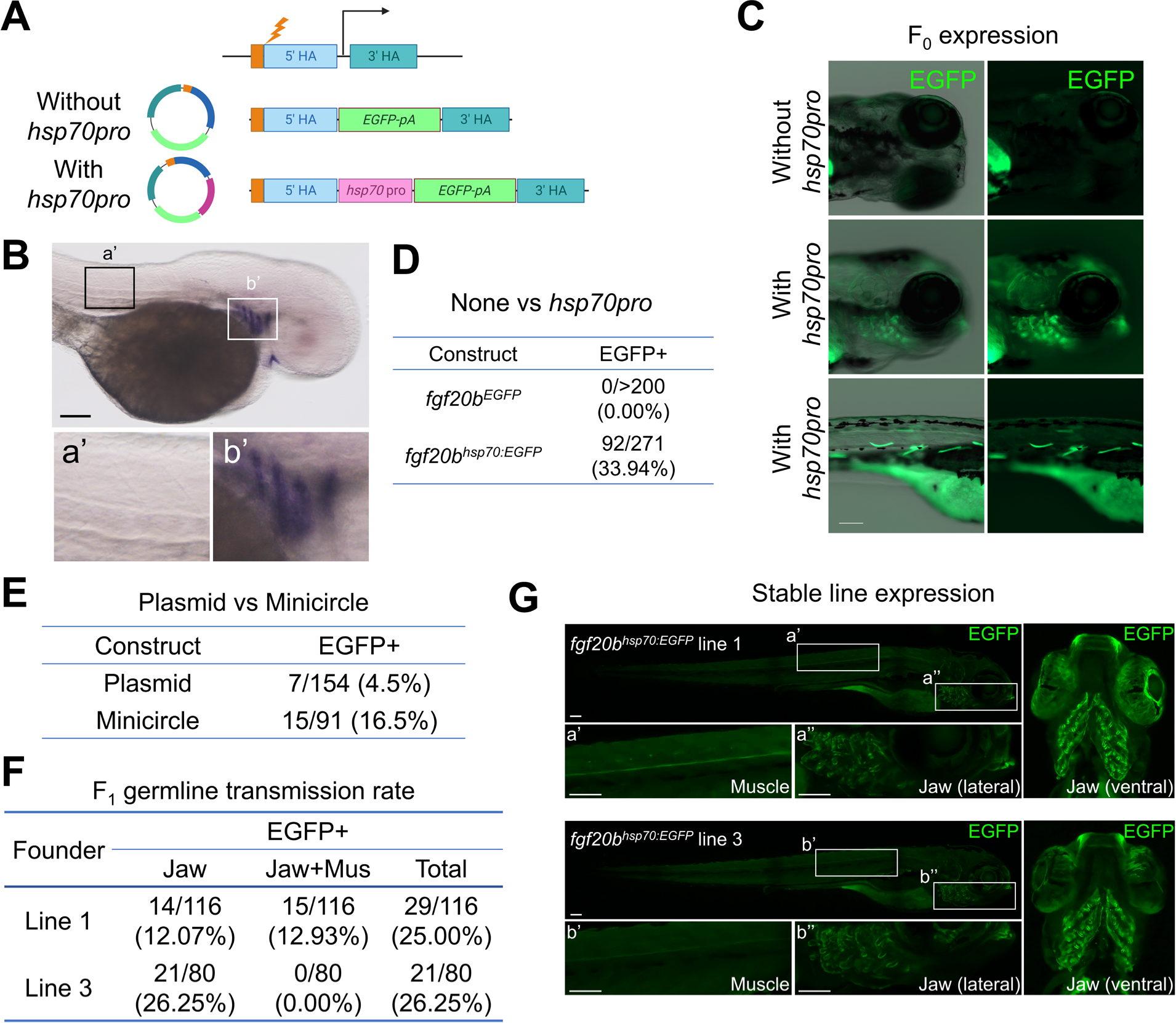
Generation of *fgf20b^hsp70:EGFP^* reporter line using a minicircle donor and *hsp70pro*. (**A**) Schematic of the strategy to integrate *EGFP* or *hsp70:EGFP* at the *fgf20b* start codon using minicircle donor. (**B**) Representative in situ hybridization (ISH) images of *fgf20b* at 1 day post-fertilization (dpf). (**C**) Representative images of F_0_ larvae injected with the *EGFP* (without *hsp70pro*) or *hsp70pro:EGFP* (with *hsp70pro*) minicircle donor. (**D**) Comparison of EGFP expressing larvae after injecting *EGFP* vs *hsp70pro:EGFP* minicircle donor. (**E**) Comparison of EGFP expressing larvae after injecting *hsp70pro:EGFP* plasmid or minicircle donor. Note that *hsp70pro:EGFP* plasmids contain two sgRNAs juxtaposed to the 5’ and 3’ HAs. (**F**) Germline transmission rate of two established *fgf20b^hsp70:EGFP^* lines. (**F**) Representative images *fgf20b^hsp70:EGFP^* stable lines. (Top) *fgf20b^hsp70:EGFP^* line 1. EGFP is observed in jaw with a restricted number of skeletal muscle cells. (Bottom) *fgf20b^hsp70:EGFP^* line 3. EGFP is detectable only in jaw. Scale bars, 100µm.

**Figure 5.**
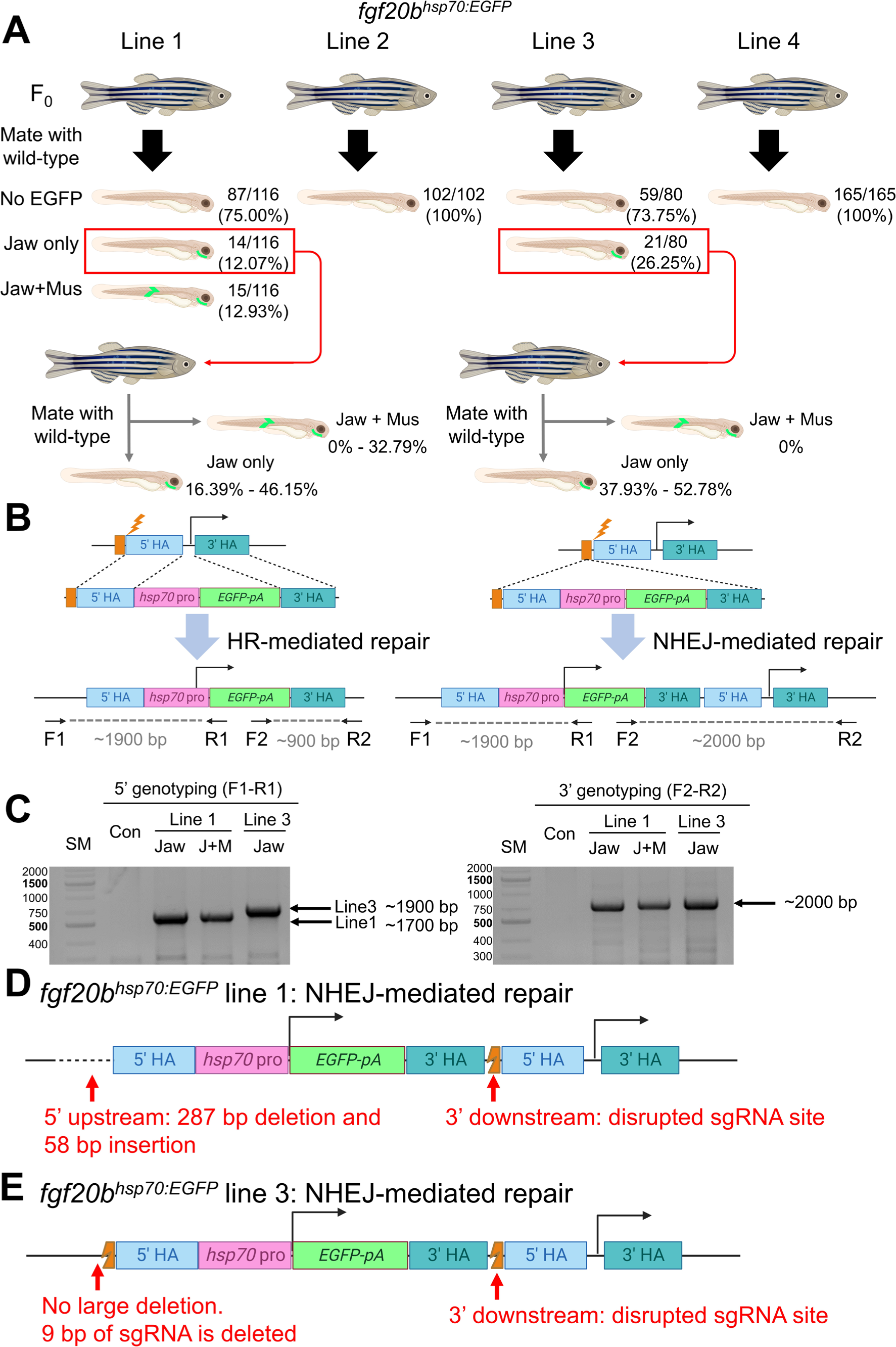
Non-specific expression of *hsp70pro* is caused by unexpected editing of the target site. (**A**) Schematic to generate *fgf20b^hsp70EGFP^* stable lines. (**B**) Schematic of KI outcome via homologous recombination (HR)-mediated repair and non-homologous end joining (NHEJ)-mediated repair mechanisms. (**C**) PCR analysis to genotype upstream and downstream regions of the integrated site at the *fgf20b* locus. Corresponding primers are distally located to the HA as shown in **(B)**. Arrows indicate correctly amplified fragment. (**D**) Schematic of the edited genomic DNA sequence for *fgf20b^hsp70:EGFP^* line 1 (top) and 3 (bottom).

We noticed that *fgf20b^hsp70:EGFP^* Line 1 produced larvae showing non-specific muscular EGFP expression while no larvae from Line 3 displayed muscular expression (**Fig. 4G, 5A, and Table S1**). Similar to *scn8ab*, the muscle expression of Line 1 displayed variation without a consistent level of muscular induction. By assessing muscular EGFP induction of progenies from the fish having jaw-specific expression, we confirmed that non-specific muscle induction in Line 1 is stochastically determined during embryogenesis (**Fig. 5A**). We next explored the causative factor behind the variation in non-specific muscular expression between *fgf20b^hsp70:EGFP^* Line 1 and Line 3 by evaluating the edited genome. PCR analysis of the 5’ upstream region revealed that Line 1 generated a slightly smaller fragment than expected, whereas Line 3 had the expected size (**Fig. 5B and C**). Sequencing identified that Line 1 had a 287-bp deletion and a 58-bp insertion at the 5’ junction (**Fig. 5D and Fig. S7**), while only the sgRNA site was disrupted in Line 3 (**Fig. 5E and Fig. S7**). Since the sgRNA site was located at the 5’ end of 5’ HA, a NHEJ-mediated KI event would lead to the presence of 5’ and 3’ HAs downstream of the inserted cassette (**Fig. 5B**). Our PCR and sequencing analyses for the 3’ downstream region confirmed the long PCR fragment, an indication of NHEJ-mediated integration (**Fig. 5B and C**). Indeed, this fragment included two copies of the 3’ HA with one 5’ HA in the middle (**Fig. 5D and E and Fig. S7**). In *fgf20b^hsp70:EGFP^* Line 3, the *EGFP* start codon of the *hsp70pro:EGFP* cassette mimicked the endogenous *fgf20b* start codon as both 5’ and 3’ HA regions were preserved. However, the 5’ upstream region of *fgf20b^hsp70:EGFP^* Line 1 had an unexpected deletion, which may contain an important regulatory element to repress *hsp70pro* leaky induction in muscular tissues. Overall, our data indicate that imperfect genome editing at the target site may influence leaky activation of the ectopic promoter.

## Discussion

Here, we report a robust KI approach in zebrafish utilizing a minicircle and a minimal promoter. Our initial attempts to integrate an *EGFP* gene at the start codon of *scn8ab* and *fgf20b* were unsuccessful in driving reporter gene expression. We speculated potential reasons for this outcome. First, the targeted genes may be transcribed at low levels below the threshold of detection, which would hinder visualization of the EGFP signal. However, this explanation is unlikely since antibody staining was unable to amplify the EGFP signal. It is plausible that integrating a long DNA fragment, such as *EGFP-pA*, may change the overall chromatin structure of the promoter region and thus subsequently influence transcription activities. Another possibility is the presence of pseudo ATGs in the 5’ untranslated region (UTR), which could serve as the TIS in the KI reporter line. For instance, the *scn8ab* transcript contains five ATGs in the 5’ UTR before the genuine start codon (**Fig. S2**). Nonetheless, our study demonstrated that placing an ectopic minimal promoter, such as *hsp70pro*, can overcome these potential problems. Ectopic promoters typically exhibit higher transcription activity, enabling the amplification of low transcription levels. Additionally, they act as a strong promoter to initiate transcription under the control of the target gene’s regulatory elements (**Fig. S8**), resolving unforeseeable transcription and translation problems. Moreover, in-frame editing is not required when using the ectopic promoters, as the inserted gene is transcribed under the control of the ectopic promoter rather than the endogenous one. Thus, incorporating an ectopic promoter provides valuable troubleshooting for creating KI lines, especially when targeting the start codon fails to direct detectable expression of the reporter gene.

The use of an ectopic promoter requires several considerations. First, the ectopic promoter may result in non-specific expression, which could potentially obscure the genuine expression of the target gene. As shown in our study, it could direct non-specific expression, mostly in skeletal muscle cells. This feature of *hsp70pro* was also reported in other studies, where *hsp70pro*-mediated enhancer trap lines displayed weak expression in heart, skeletal muscle, and lens (Asakawa et al., 2008; Kimura et al., 2014; Nagayoshi et al., 2008). Addressing the issue of non-specific expression, which is highly restricted to a few skeletal muscle cells in most larvae, can be accomplished by meticulously sorting fish with minimal non-specific expression. We also found that non-specific expression can be induced when the edited genome is significantly modified by deletion. When the integration of a donor construct causes minimal sequence change, non-specific expression is undetectable as shown in the case of *fgf20b^hsp70:EGFP^* Line 3. These results suggest that the undesired activation of the ectopic promoter can be repressed by preserving the upstream and downstream sequences in the edited genome. Therefore, careful evaluation of the potential off-target effects should be conducted. For example, establishing the most plausible line would require comparing the expression patterns of multiple independent lines. Another consideration is that the established line may not accurately recapitulate the endogenous regulation of the target gene, as the expression level may not reflect the true physiological conditions. Further assessment of a minimal promoter lacking basal activity, such as the mouse beta-globin minimal promoter {Kemmler, 2023 #4876} or the zebrafish synthetic promoter {Begeman, 2022 #4831}, will be essential to strengthen our new KI approach.

We assess the KI strategy using a minicircle as a donor dsDNA. Under our experimental conditions, a minicircle can highly enhance the KI efficiency compared to a plasmid. A similar finding was also reported in post-mitotic cells of the rat model (Suzuki et al., 2016). Previous studies reported that internal linearization via injecting a donor plasmid that harbors cleavage sites appears to be more efficient than externally linearized dsDNA via PCR or enzyme digestion (Carroll and Beumer, 2014; Hoshijima et al., 2016; Irion et al., 2014). Following these reports, a minicircle can be an excellent circular form of donor DNA. In addition, linearization of a minicircle can be achieved by one sgRNA site, one less than that of a plasmid which typically requires two sgRNAs at the juxtaposition of the HAs (Hoshijima et al., 2016). Two sgRNAs in a plasmid are necessary to avoid the integration of unnecessary elements, such as antibiotic-resistant genes and the origin of replication, when integration occurs via NHEJ (Auer et al., 2014; Kesavan et al., 2017). The enhanced efficiency of the minicircle can be explained by its short length, lack of bacterial sequences, and minimal number of linearization sites. In summary, our study demonstrates that a minicircle serves as a powerful donor DNA for the KI strategy.

Our strategy employs distinct choice for sgRNAs targeting *scn8ab* and *fgf20b*. While *scn8ab* sgRNA is positioned between 5’ and 3’ HAs, *fgf20b* sgRNA is placed at the 5’ end of 5’ HA. Notably, all *fgf20b* KI lines result from NHEJ-mediated integration, thereby having two copies of both 5’ and 3’ HAs in the genome. Similar outcomes were also reported from other studies {Auer, 2014 #4870}{Kesavan, 2017 #4863}, where sgRNAs were located at the end of or within the middle of 5’ HAs. Moreover, another study revealed that donor vectors lacking HAs can trigger NHEJ-mediated integration {Li, 2019 #5230}. Although more data require to establish conclusive findings, these observations suggest that the absence of a donor template connecting the double strand break (DSB) at the cleavage site may induce NHEJ-mediated integration rather than HR-mediated repair. Thus, sgRNA position would be considered for more precise designing of KI generation.

Overall, our strategy provides an efficient method to generate KI reporter lines, expanding our capacity to create KI animals and further facilitating the study of gene expression impacting diverse biological events.

## Acknowledgments

We thank the UW–Madison SMPH BRMS staff and members of the Kang Lab for zebrafish care; Tamplin lab for technical support; and Chen-Hui Chen, Jingli Cao, and Kang lab members for comments on the manuscript.

## Author Contributions

Biological Experiments: MK, RH, DO, ARP, SIA, JK

Conceptualization: MK, RH, DO, JK

Writing, Reviewing, Editing: MK, RH, DO, JK

Funding: RH, DO, JK

