## Supplementary information for "A robust knock-in approach using a minimal promoter and a minicircle"

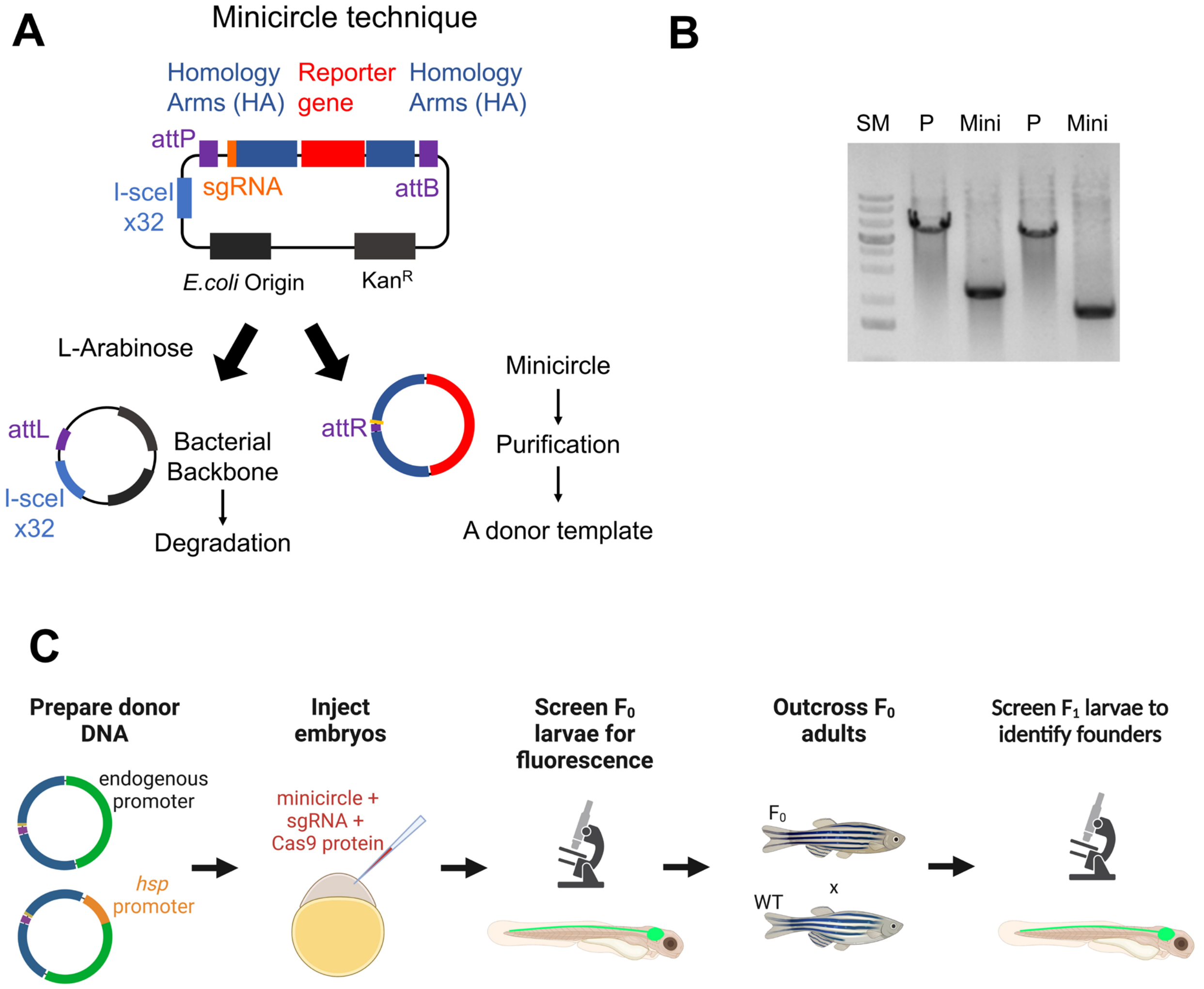


**Figure S1. Schematic strategy to generate KI reporter line using minicircle.** (**A**) Overview of minicircle generation. (**B**) A representative gel picture of parental plasmids and minicircles. SM, size marker. P, parental plasmid. Mini, minicircle. (**C**) Schematic of KI generation strategy.

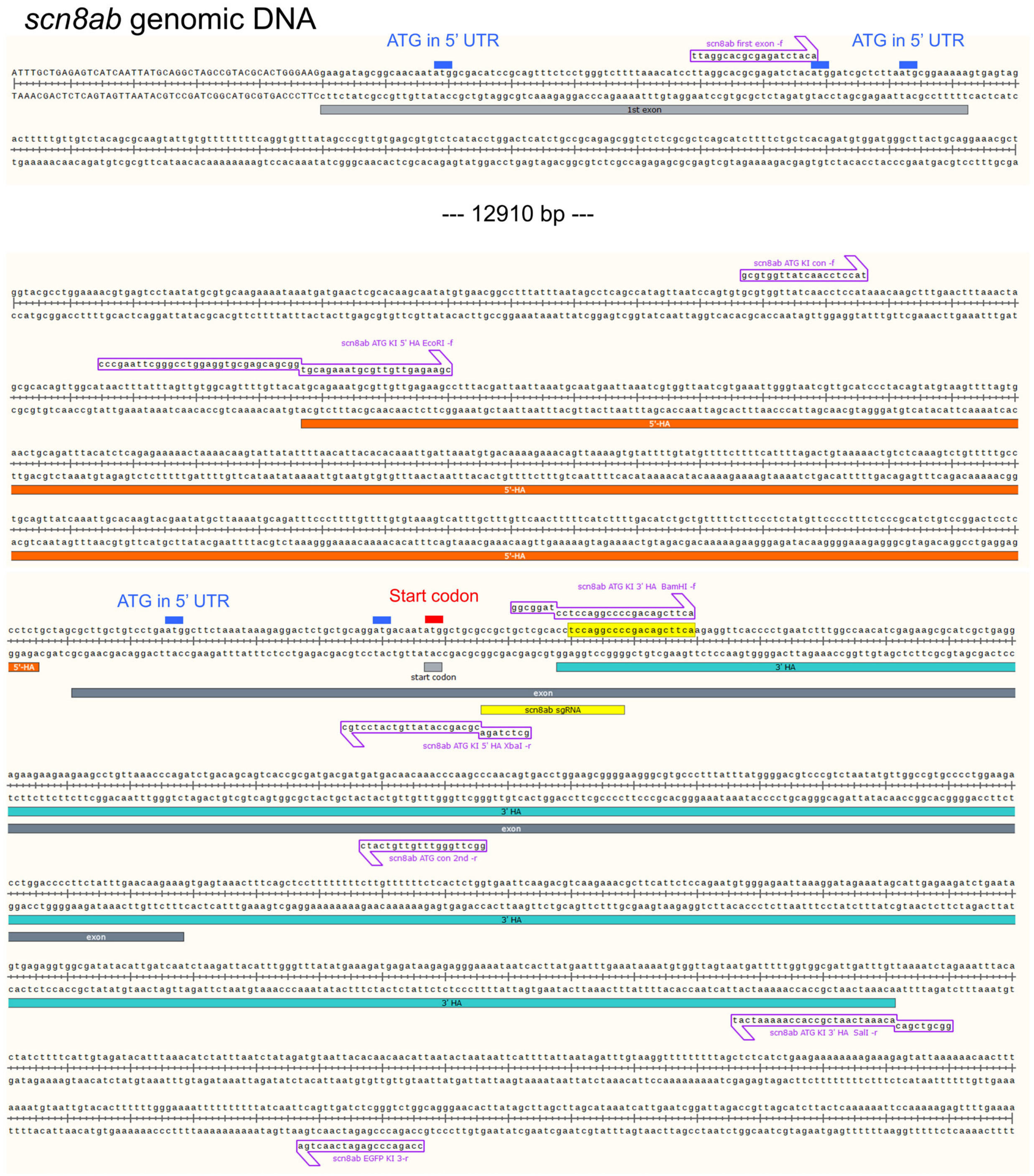


**Figure S2. Genomic DNA sequence of *scn8ab*.** The first and second exons of *scn8ab* are separated by a distance of 13.7kb. Blue bars indicate five pseudo-ATG start codon in 5' untranslated region (UTR). Red bar indicates *scn8ab* start codon.

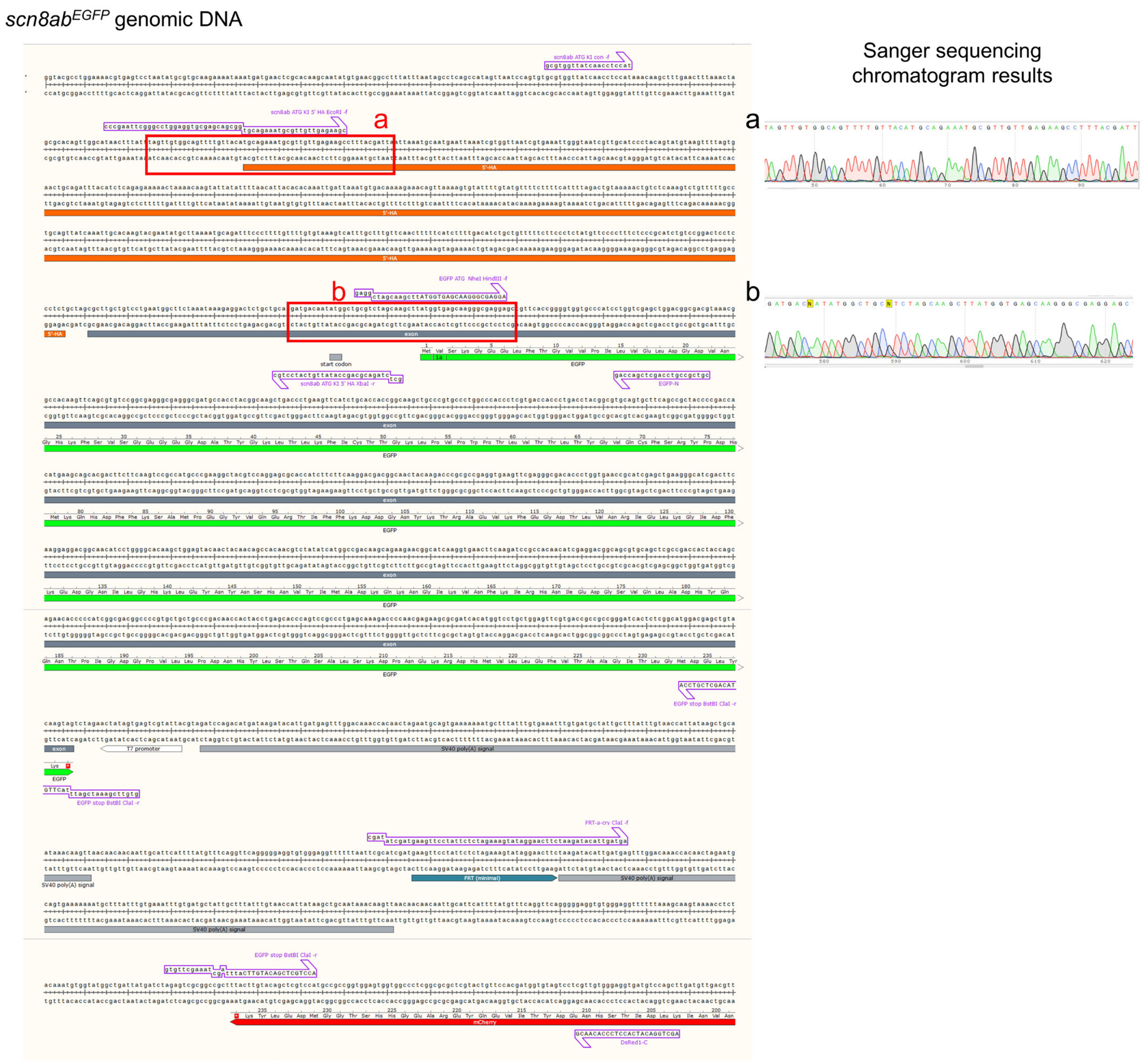
scn8ab$^{EGFP}$ genomic DNA
Sanger sequencing chromatogram results
a
b

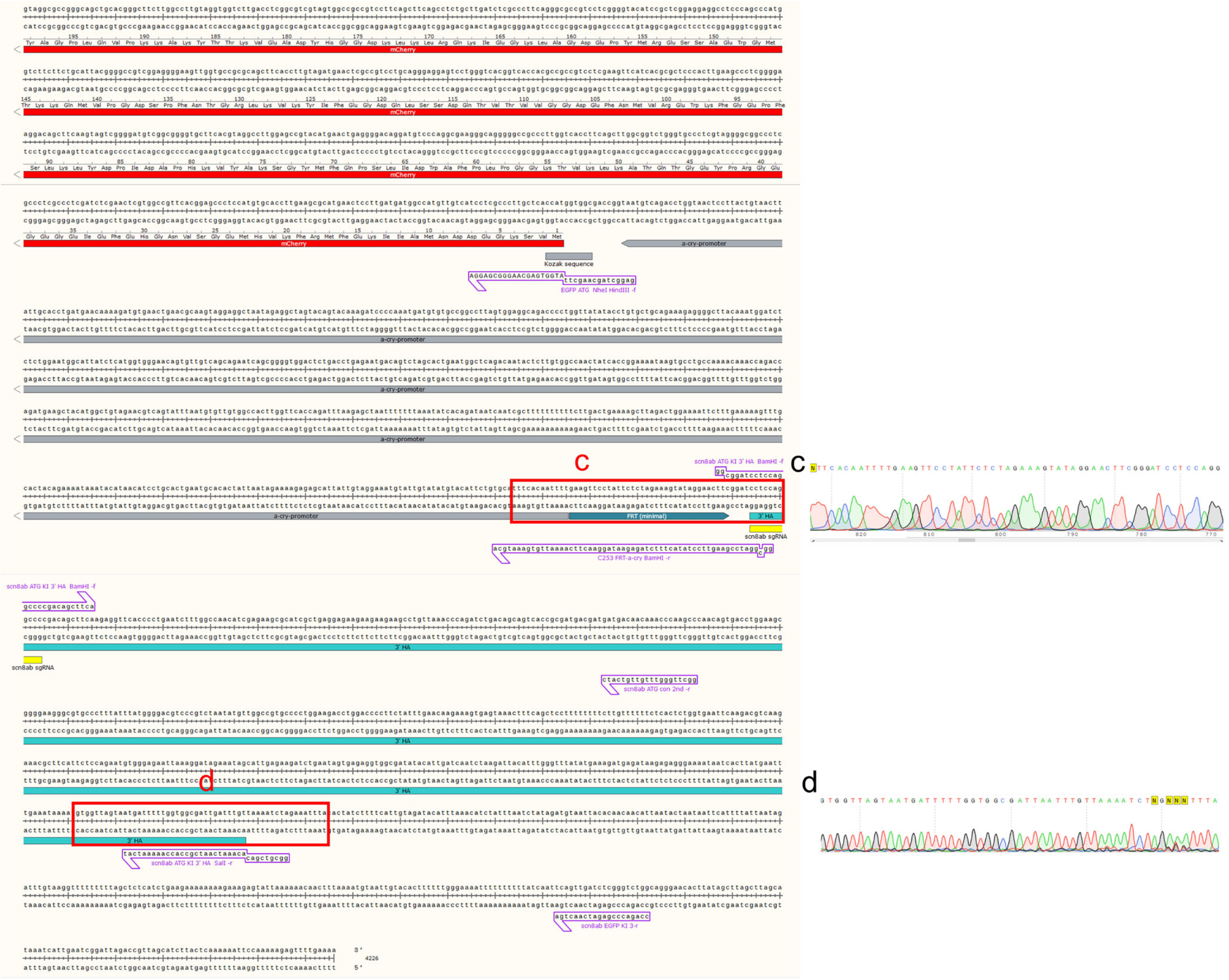

**Figure S3. Genomic DNA sequence of *scn8ab*$^{EGFP}$.** Sanger sequencing chromatogram results are shown in the right.

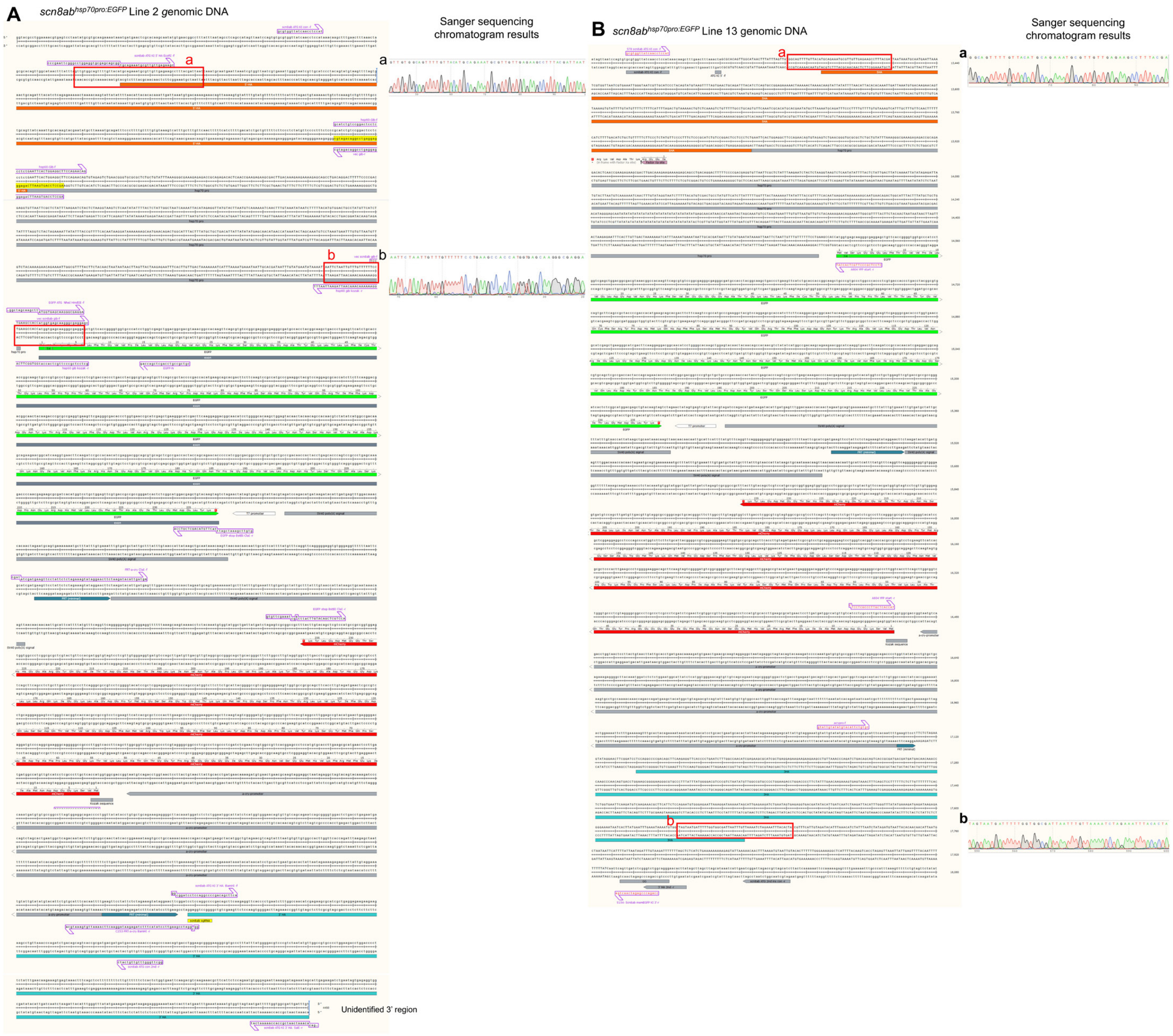
A
scn8ab hsp70pro:EGFP Line 2 genomic DNA
Sanger sequencing chromatogram results
a
b
Unidentified 3' region
B
scn8ab hsp70pro:EGFP Line 13 genomic DNA
Sanger sequencing chromatogram results
a
b

**Figure S4. Genomic DNA sequence of *scn8ab*$^{hsp70:EGFP}$ line 2 (A) and 13 (B).** Sanger sequencing chromatogram results are shown in the right.

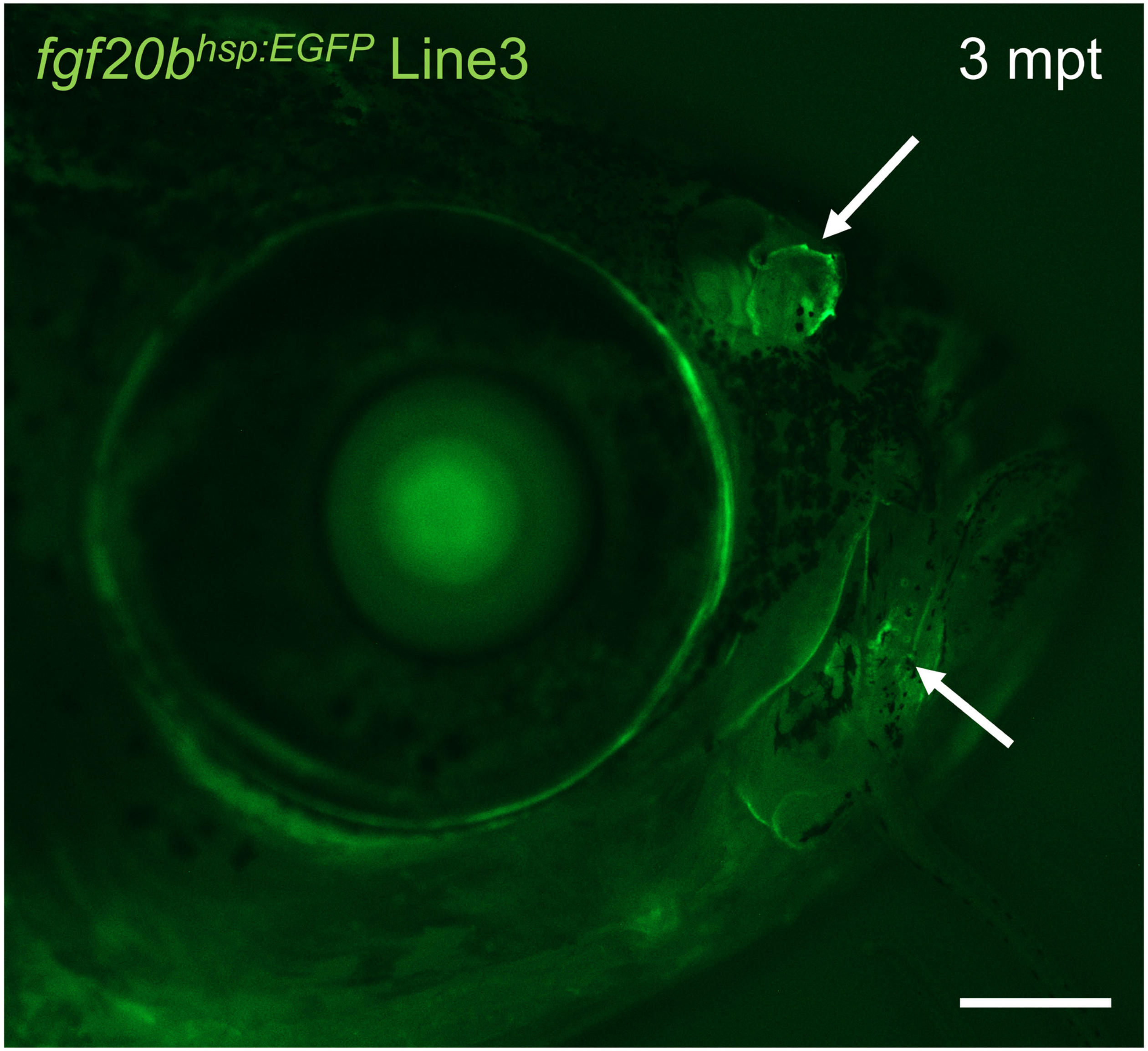


**Fig. S5. A representative image of *fgf20b*$^{hspEGFP}$ line 3 at 3 mpt**

### *fgf20b* genomic DNA

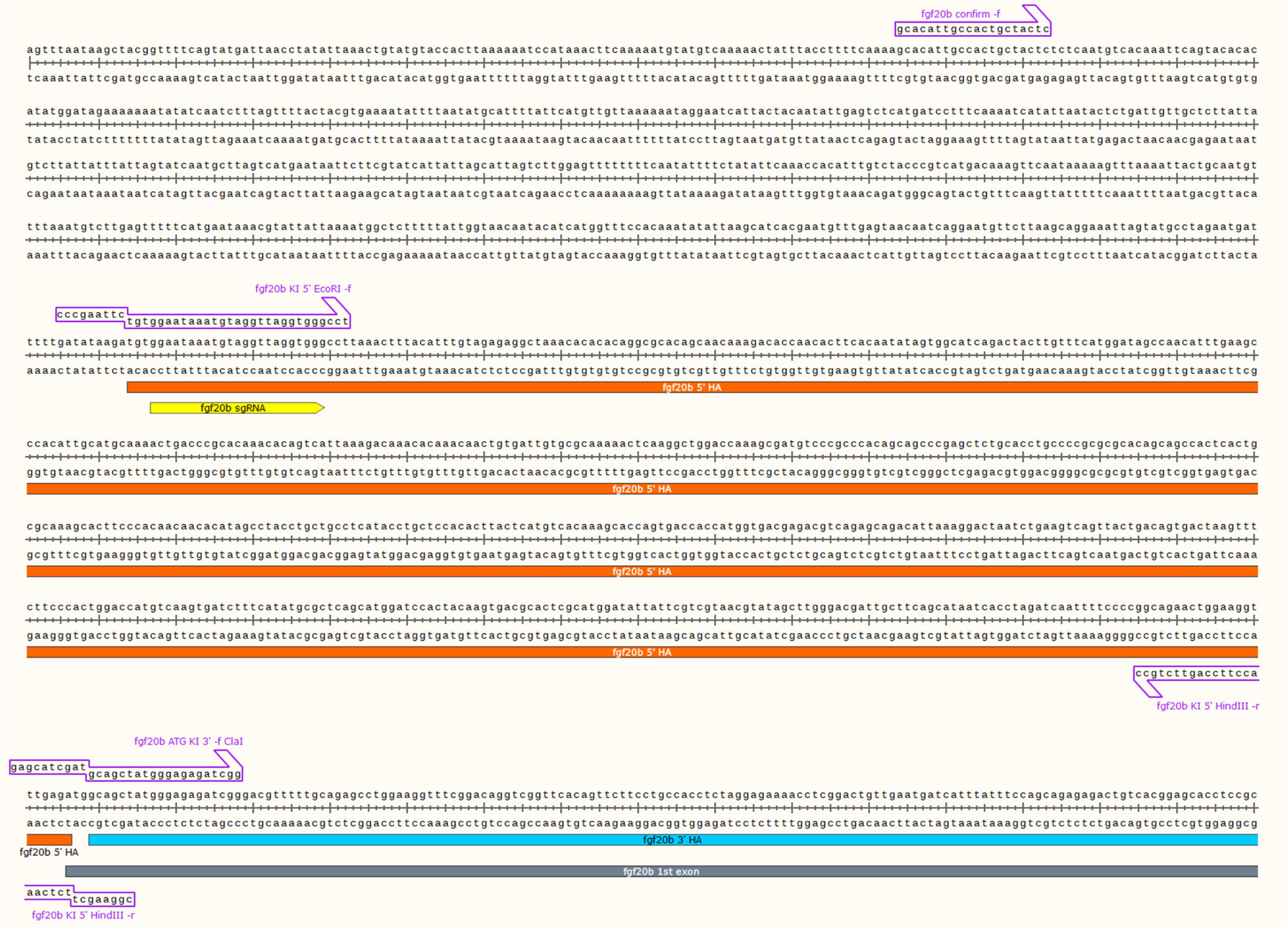

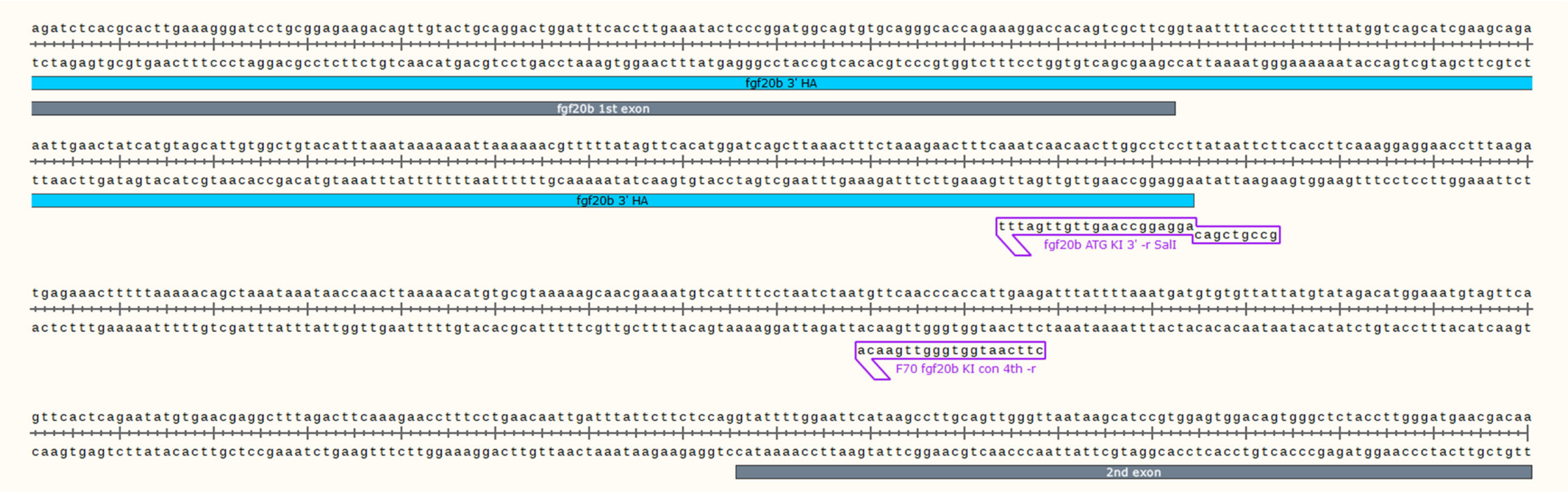


**Figure S6. Genomic DNA sequence of *fgf20b*.**

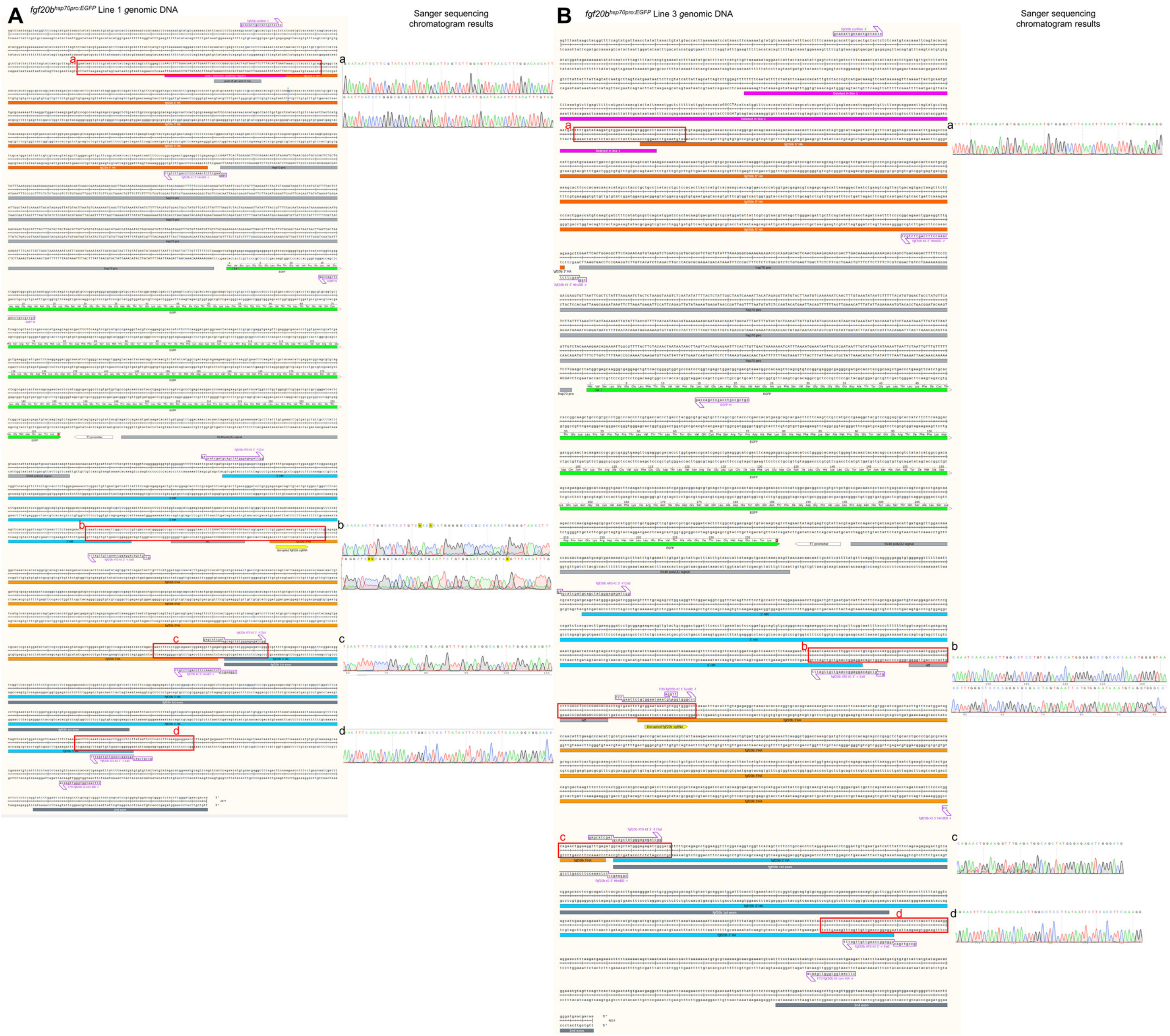
A
fgf20b^hsp70pro:EGFP Line 1 genomic DNA
Sanger sequencing chromatogram results
B
fgf20b^hsp70pro:EGFP Line 3 genomic DNA
Sanger sequencing chromatogram results

**Figure S7. Genomic DNA sequence of *fgf20b*$^{hsp70:EGFP}$ line 1 (A) and 3 (B).** Sanger sequencing chromatogram results are shown in the right.

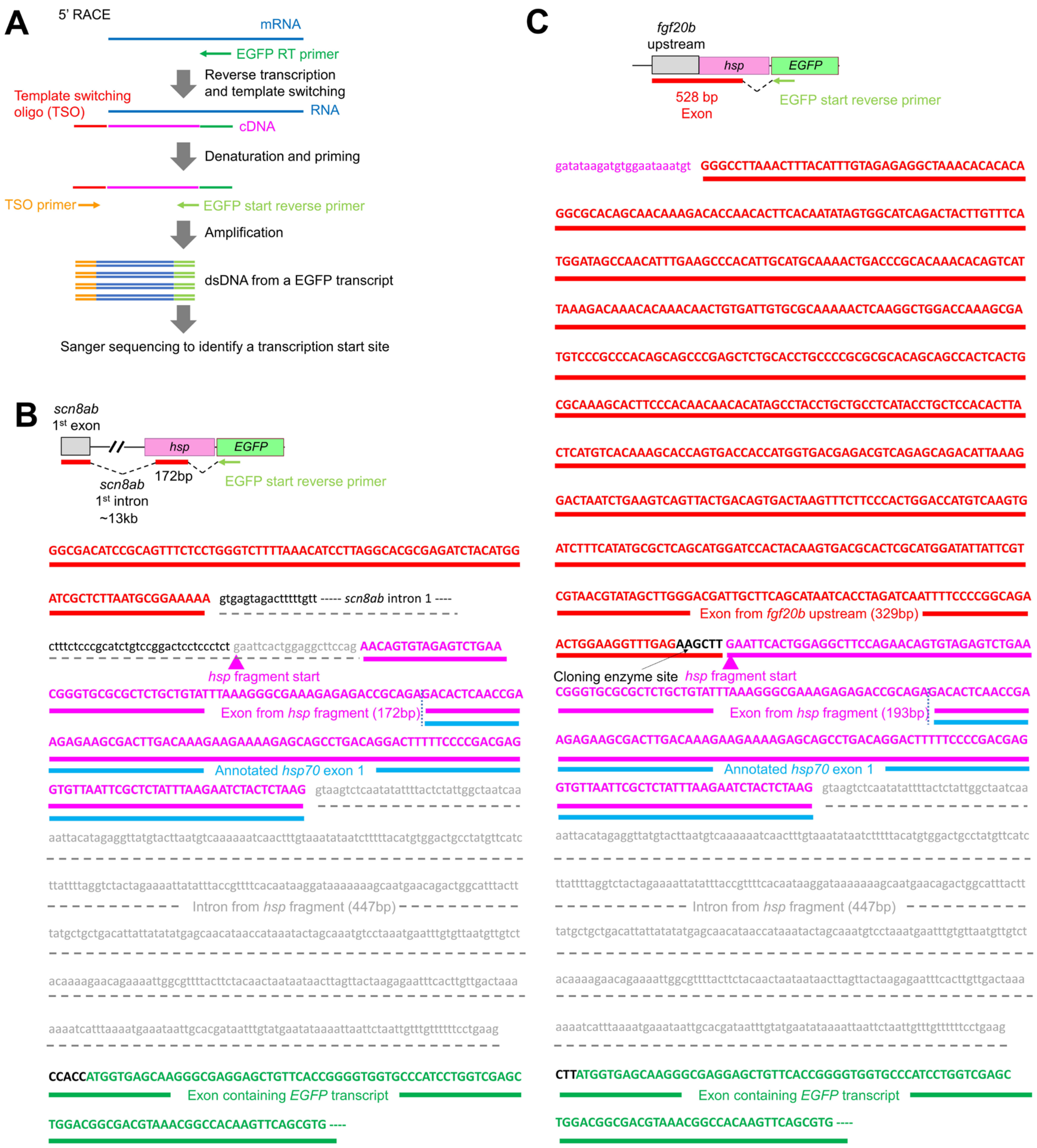
A
5' RACE
mRNA
EGFP RT primer
Reverse transcription
and template switching
Template switching
oligo (TSO)
RNA
cDNA
Denaturation and priming
TSO primer
EGFP start reverse primer
Amplification
dsDNA from a EGFP transcript
Sanger sequencing to identify a transcription start site
B
scn8ab
1st exon
hsp
EGFP
172bp
scn8ab
1st intron
~13kb
EGFP start reverse primer
GGCGACATCCGCAGTTTCTCCTGGGTCTTTTAAACATCCTTAGGCACGCGAGATCTACATGG
ATCGCTCTTAATGCGGAAAAA gtgagtagactttttgtt ----- scn8ab intron 1 ----
ctttctcccgcatctgtccggactcctccctct gaattcactggaggcttccag AACAGTGTAGAGTCTGAA
hsp fragment start
CGGGTGCGCGCTCTGCTGTATTTAAAGGGCGAAAGAGAGACCGCAGAGACACTCAACCGA
Exon from hsp fragment (172bp)
AGAGAAGCGACTTGACAAAGAAGAAAAGAGCAGCCTGACAGGACTTTTTCCCCGACGAG
Annotated hsp70 exon 1
GTGTTAATTCGCTCTATTTAAGAATCTACTCTAAG gtaagtctcaatatattttactctattggctaatcaa
aattacatagaggttatgtacttaatgtcaaaaaatcaactttgtaaatataatctttttacatgtggactgcctatgttcatc
ttattttaggtctactagaaaattatatttaccgttttcacaataaggataaaaaaagcaatgaacagactggcatttactt
Intron from hsp fragment (447bp)
tatgctgctgacattattatatatgagcaacataaccataaatactagcaaatgtcctaaatgaatttgtgttaatgttgtct
acaaaagaacagaaaattggcgttttacttctacaactaataataacttagttactaagagaatttcacttgttgactaaa
aaaatcatttaaaatgaaataattgcacgataatttgtatgaatataaaattaattctaattgtttgtttttttcctgaag
CCACCATGGTGAGCAAGGGCGAGGAGCTGTTCACCGGGGTGGTGCCCATCCTGGTCGAGC
Exon containing EGFP transcript
TGGACGGCGACGTAAACGGCCACAAGTTCAGCGTG ----
C
fgf20b
upstream
hsp
EGFP
528 bp
Exon
EGFP start reverse primer
gatataagatgtggaataaatgt GGGCCTTAAACTTTACATTTGTAGAGAGGCTAAACACACACA
GGCGCACAGCAACAAAGACACCAACACTTCACAATATAGTGGCATCAGACTACTTGTTTCA
TGGATAGCCAACATTTGAAGCCCACATTGCATGCAAAACTGACCCGCACAAACACAGTCAT
TAAAGACAAACACAAACAACTGTGATTGTGCGCAAAAACTCAAGGCTGGACCAAAGCGA
TGTCCCGCCCACAGCAGCCCGAGCTCTGCACCTGCCCCGCGCGCACAGCAGCCACTCACTG
CGCAAAGCACTTCCCACAACAACACATAGCCTACCTGCTGCCTCATACCTGCTCCACACTTA
CTCATGTCACAAAGCACCAGTGACCACCATGGTGACGAGACGTCAGAGCAGACATTAAAG
GACTAATCTGAAGTCAGTTACTGACAGTGACTAAGTTTCTTCCCACTGGACCATGTCAAGTG
ATCTTTCATATGCGCTCAGCATGGATCCACTACAAGTGACGCACTCGCATGGATATTATTCGT
CGTAACGTATAGCTTGGGACGATTGCTTCAGCATAATCACCTAGATCAATTTTCCCCGGCAGA
Exon from fgf20b upstream (329bp)
ACTGGAAGGTTTGAGAAGCTT GAATTCACTGGAGGCTTCCAGAACAGTGTAGAGTCTGAA
Cloning enzyme site
hsp fragment start
CGGGTGCGCGCTCTGCTGTATTTAAAGGGCGAAAGAGAGACCGCAGAGACACTCAACCGA
Exon from hsp fragment (193bp)
AGAGAAGCGACTTGACAAAGAAGAAAAGAGCAGCCTGACAGGACTTTTTCCCCGACGAG
Annotated hsp70 exon 1
GTGTTAATTCGCTCTATTTAAGAATCTACTCTAAG gtaagtctcaatatattttactctattggctaatcaa
aattacatagaggttatgtacttaatgtcaaaaaatcaactttgtaaatataatctttttacatgtggactgcctatgttcatc
ttattttaggtctactagaaaattatatttaccgttttcacaataaggataaaaaaagcaatgaacagactggcatttactt
Intron from hsp fragment (447bp)
tatgctgctgacattattatatatgagcaacataaccataaatactagcaaatgtcctaaatgaatttgtgttaatgttgtct
acaaaagaacagaaaattggcgttttacttctacaactaataataacttagttactaagagaatttcacttgttgactaaa
aaaatcatttaaaatgaaataattgcacgataatttgtatgaatataaaattaattctaattgtttgtttttttcctgaag
CTTATGGTGAGCAAGGGCGAGGAGCTGTTCACCGGGGTGGTGCCCATCCTGGTCGAGC
Exon containing EGFP transcript
TGGACGGCGACGTAAACGGCCACAAGTTCAGCGTG ----

**Fig. S8. 5' RACE results of *scn8ab*$^{hspEGFP}$ line 2 and *fgf20b*$^{hspEGFP}$ line 3. (A)** The schematic of 5' RACE experiment to identify the transcription start site. (**B**) 5' RACE result of *scn8ab*$^{hspEGFP}$ line 2. Solid lines indicate the exon sequenced by the RACE experiment. Dotted lines represent introns. Transcription for *scn8ab*$^{hspEGFP}$ appears to initiate from the endogenous exon 1 of *scn8ab* (Red bar, Red uppercase), which is located ~13kb away from the integration site. The 5' region of the *hsp* fragment serves as a new splicing acceptor, resulting in the transcription of a 172-bp fragment (2$^{nd}$ exon). Note that the first exon of *hsp70* gene (blue line) is a part of this fragment. Another splicing event occurs within the *hsp70* fragment, and the third exon starts near the EGFP gene. (**C**) 5' RACE result of *fgf20b*$^{hspEGFP}$ line 3. Solid lines indicate the exon sequenced by the RACE experiment. Dotted lines represent introns. Transcription for *fgf20b*$^{hspEGFP}$ appears to initiate from the upstream region of *fgf20b* (Red bar, Red uppercase). The first exon extends into the 5' region of *hsp* fragment, resulting in the transcription of 528-bp. Note that the first exon of *hsp70* gene (blue line) is a part of this fragment. Another splicing event occurs within the *hsp70* fragment, and the third exon starts near the EGFP gene.

**Supplementary Table**

**Table S1. Mating results**

(**A**) Mating results of *scn8ab*$^{hsp70:EGFP/+}$ line 2 and wild-type. Expression is determined with larvae.

*scn8ab*$^{hsp70:EGFP/+}$ line 2 and wild-type

| | Neuron only | Neu + Mus (weak) | Neu + Mus (moderate) | Neu + Mus (strong) | No expression | Total |
|---|---|---|---|---|---|---|
| Pair 1 | 57 (28.5%) | 31 (15.5%) | 5 (2.5%) | 1 (0.5%) | 106 (53.5%) | 200 |
| Pair 2 | 0 (0.0%) | 10 (29.4%) | 5 (14.7%) | 0 (0.0%) | 19 (55.9%) | 34 |
| Pair 3 | 5 (5.3%) | 16 (16.8%) | 7 (7.4%) | 10 (10.5%) | 57 (60.0%) | 95 |
| Pair 4 | 2 (4.3%) | 15 (31.9%) | 5 (10.6%) | 0 (0.0%) | 25 (53.2%) | 47 |
| Pair 5 | 2 (6.7%) | 11 (36.7%) | 2 (6.7%) | 0 (0.0%) | 15 (50.5%) | 30 |
| Pair 6 | 6 (2.8%) | 124 (57.7%) | 0 (0.0%) | 0 (0.0%) | 85 (39.5%) | 215 |
| Pair 7 | 13 (11.8%) | 24 (21.8%) | 11 (10.0%) | 3 (2.7%) | 59 (53.6%) | 110 |
| Pair 8 | 9 (8.6%) | 48 (45.7%) | 0 (0.0%) | 0 (0.0%) | 48 (45.7%) | 105 |
| Pair 9 | 16 (18.4%) | 35 (40.2%) | 0 (0.0%) | 0 (0.0%) | 36 (41.4%) | 87 |

(**B**) Mating results of *fgf20b*$^{hsp70:EGFP/+}$ line 1 and wild-type. Expression is determined with larvae.

*fgf20b*$^{hsp70:EGFP/+}$ line 1 and wild-type

| | Jaw only EGFP | Jaw + Mus | No expression | Total |
|---|---|---|---|---|
| Pair 1 | 36 (46.15%) | 0 (0%) | 42 (53.8%) | 78 |
| Pair 2 | 24 (42.10%) | 8 (14.04%) | 25 (43.86%) | 57 |
| Pair 3 | 15 (28.84%) | 12 (23.08%) | 25 (48.08%) | 52 |
| Pair 4 | 10 (16.39%) | 20 (32.79%) | 31 (50.82%) | 61 |
| Pair 5 | 2 (11.11%) | 5 (27.78%) | 11 (61.11%) | 18 |
| Pair 6 | 13 (24.53%) | 10 (18.87%) | 30 (56.60%) | 53 |
| Pair 7 | 5 (62.5%) | 0 (0%) | 3 (37.5%) | 8 |
| Pair 8 | 23 (38.33%) | 4 (6.67%) | 33 (55%) | 60 |

(**C**) Mating results of *fgf20b*$^{hsp70:EGFP/+}$ line 3 and wild-type. Expression is determined with larvae.

| *fgf20b*$^{hsp70:EGFP/+}$ line 3 and wild-type | | | | |
|---|---|---|---|---|
| | Jaw only EGFP | Jaw + Mus | No expression | Total |
| Pair 1 | 19 (52.78%) | 0 (0.00%) | 17 (47.22%) | 36 |
| Pair 2 | 11 (37.93%) | 0 (0.00%) | 18 (62.07%) | 29 |
| Pair 3 | 16 (45.71%) | 0 (0.00%) | 19 (54.29%) | 35 |
| Pair 4 | 11 (52.38%) | 0 (0.00%) | 10 (47.62%) | 21 |

**Table S2. Primers used in this study**

| | |
|---|---|
| ***scn8ab* genotyping** | |
| *5' junction* | |
| scn8ab ATG KI con -f | gcgtggttatcaacctccat |
| EGFP-N | CGTCGCCGTCCAGCTCGACCAG |
| *3' junction* | |
| Acrypro-F from Ryan | Gtattgtatatgtacattctgtgc |
| scn8ab EGFP KI 3-r | ccagacccgagatcaactga |
| *scn8ab RT-PCR* | |
| scn8ab first exon -f | ttaggcacgcgagatctaca |
| scn8ab ATG con 2nd -r | ggcttgggtttgttgtcatc |
| EGFP-N | CGTCGCCGTCCAGCTCGACCAG |
| ***fgf20b* genotyping** | |
| *5' junction* | |
| fgf20b confirm -f | gcacattgccactgctactc |

| | |
|---|---|
| hsp70 pro HindIII -r | ggt aagctt CAGGAAAAACAAACAATTAGAATTAATTT |
| *3' junction* | |
| GFP confirm -f | AGAAGCGCGATCACATGGTC |
| fgf20b KI confirm 4th -r | cttcaatggtgggttgaaca |
| **in situ hybridization** | |
| fgf20b ATG MseI -f | ccc caattg atggcagctatgggagagat |
| fgf20b stop XbaI -r | cc tctaga tca actgtgtccgagcacct |
| scn8ab ISH 2nd -f | ACGTCACTGGCGTATCGAG |
| scn8ab ISH 2nd -r | CATGGGCAAAATGTTGAATG |
| ***scn8ab* plasmids** | |
| EGFP ATG NheI HindIII -f | gag gctagc aagctt ATG GTG AGC AAG GGC GAG GA |
| EGFP stop BstBI ClaI -r | gtg ttcgaa atcgat ttaCTTGTACAGCTCGTCCA |
| FRT-a-cry ClaI -f | cgat atcgat gaagttcctattctctagaaagtataggaacttc taagatacattgatga |
| FRT-a-cry BamHI -r | ggc ggatcc gaagttcctatactttctagagaataggaacttcaaaattgtgaaatgca |
| scn8ab ATG KI 5' HA EcoRI -f | ccc gaattc gggcctggaggtgcgagcag cgg tgcagaaatgcgttgttgagaagc |
| scn8ab ATG KI 5' HA XbaI -r | gc tctaga cgcagccatattgtcatcctgc |
| scn8ab ATG KI 3' HA BamHI -f | ggc ggatcc tccaggccccgacagcttca |
| scn8ab ATG KI 3' HA SalI -r | ggc gtcgac acaaatcaatcgccaccaaaaatcat |
| Hsp60-gib-f | gcatctgtccggactcctccctctgaattcactggaggcttccagaacag |
| hsp60 gib kozak-r | gctcctcgcccttgctcaccatggtggcttcaggaaaaacaaacaattagaattaattt |
| vec scn8ab gib-f | tcctgaagccaccatggtgagcaagggcgaggagc |
| vec gib-r | agcctccagtgaattcagagggaggagtccggacagatgc |
| scn8ab PP KLD-f | cctccaggcccGTCGACCCATGGGGGCCC |
| scn8ab PP KLD-r | tgcgagcagcggACAAATCAATCGCCACCAAAAATCATTACTAACCAC |

| ***fgf20b* plasmids** | |
|---|---|
| fgf20b KI 5' EcoRI -f | ccc gaattc tgtggaataaatgtaggttaggtgggcct |
| fgf20b KI 5' HindIII -r | cgg aagctt ctcaaaccttccagttctgcc |
| fgf20b ATG KI 3' -f ClaI | gagc atcgat gcagctatgggagagatcgg |
| fgf20b ATG KI 3' -r SalI | gcc gtcgac aggaggccaagttgttgattt |
| hsp70 pro HindIII -f | ggt aagctt ACTGGAGGCTTCCAGAACAGT |
| hsp70 pro HindIII -r | ggt aagctt CAGGAAAAAACAAACAATTAGAATTAATTT |
| fgf20b 5'HA CAGT BsaI -f | atc ggtctc t CAGT tgtggaataaatgtaggttaggtgggcct |
| fgf20b 5'HA AGTC BsaI -r | gat GGTCTC T GACT tctcaaaccttccagttctgcc |
| fgf20b 3'HA CTCA BsaI -f | atc GGTCTC T CTCA gcagctatgggagagatcgg |
| fgf20b 3'HA CCAA BsaI -r | gat GGTCTC T TTGG aggaggccaagttgttgattt |
| fgf20b two sgRNA 3'HA CCAA BsaI -r | gat GGTCTC T TTGG ccc accta accta cattt attcc aggaggccaagttgttgattt |

#### Supplementary Video S1.

Swimming behavior of control (Left) and compound heterozygote of *scn8ab*$^{temca/hsp70:EGFP}$ (Right) at 33 °C

#### Supplementary Method

#### Subcloning pMC donor plasmid

1. Choose your knock-in (KI) strategy
a) Targeting upstream of ATG (integrate into the promoter region)
  - Pros: Frame will not affect transcription as a gene of interest is integrated upstream of ATG.
  - Con: An ectopic promoter, such as *hsp70* minimal promoter, may be required if expression of a reporter gene is not detectable.

b) Targeting middle of exon
  - Pros: Recapitulate the endogenous expression.
  - Con: Only in-frame editing can direct expression.

2. Select sgRNA site
Two options
a) sgRNA site is located in the 5' HA; 5' HA usually starts with the sgRNA site.
b) sgRNA site is located between 5' and 3' HAs. -> This sgRNA site is located at the 5' end of 5' HA.
* Note: CRISPRscan web tool would help select a highly efficient sgRNA sequence (https://www.crisprscan.org/).

3. Subclone 5' and 3'HA into the mini-circle plasmid using restriction enzyme-based or Gibson subcloning approaches

#### Minicircle DNA purification

1. Transform the pMC vector into ZYCY10P3S2T *E.coli*.

2. Day 1 morning: Seed culture: 3ml LB containing 50ug/ml Kanamycin at 37ºC, with shaking at 250 rpm

3. Day 1 afternoon: overnight culture
   a) Prepare 50ml LK media in 500ml flask
   b) Add 1ml of seed culture and culture overnight at 37 ºC, with shaking at 250 rpm

4. Next morning, prepare a Minicircle Induction Mix
   a) Dissolve L-Arabinose to make 20% solution (i.e. 0.1g into 0.5ml distilled water)
   b) Transfer 50ml LB (without kanamycin) into the overnight cultured flask
   c) Add 200ul of 20% L-Arabinose solution into the flask

5. Incubate at 30 degree with shaking at 250 rpm for 6 hours or longer

6. Harvest *E.coli* and perform Midi-prep using a commercial kit, such as Takara Extra Midi EF Kit (cat# 740420).

   Note: After harvest cells and discard liquid media, *E.coli* can be stored in the freezer until midi prep is performed.

7. Elute with 40-50ul of RNase free water

8. Cut with enz to check minicircle vector

   First lane: parental plasmid

   Next lane: mini circle plasmid

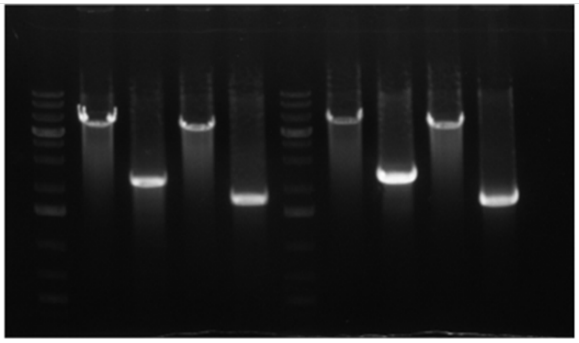

9. If there are contaminants such as *E.coli* gDNA or parental DNA, cut with 1-2 enzymes, which cut parental plasmid but not minicircle DNA. Linearized contaminants can be digested by plasmid-safe ATP-dependent DNase.
   a) Add restriction enzyme and incubate 37 degree 3 hours to overnight
   b) Treat plasmid-safe ATP-dependent DNase (Epicentre, E3101K)
      If you use a cutsmart buffer, add 1/10 10mM ATP (final 1mM), 1/10 5mM DTT (final 0.5mM), 2/25 DNase
      Note: ATP with the kit is 25mM, not 10mM.
   c) Incubate 2-16 hours 37 degree

d) (Optional) Inactivate DNase at 70 degree for 30 min
e) Purify DNA with column-based or other method
f) Elute with RNase free water
g) Cut with enzyme and check whether minicircle is intact

Examples of contamination with parental and E.coli gDNA

First lane: parental plasmid

Next lane: mini circle plasmid

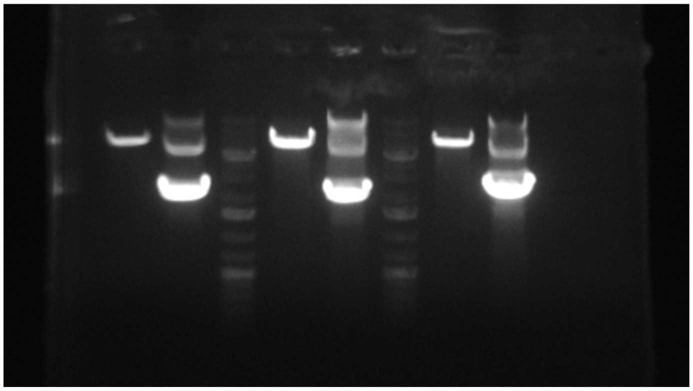

Cut with XhoI and ApaLI

plasmid-safe Dnase Treatment

Purify with column

SalI cut (for linearization)

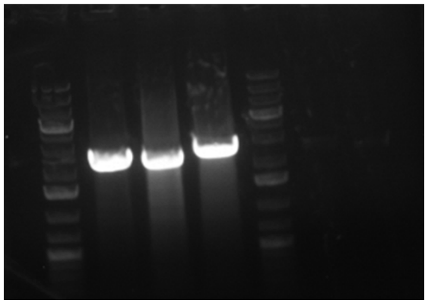

Microinjection to create KI lines

1) Injection mixture

| Component | Volume | Stock concentration | Final concentration |
|---|---|---|---|
| sgRNA | 1 ul | 120 ng/ul | 25-30 ng/ul |
| Cas9 protein | 2 ul | 1 ug/ul | 0.5 ug/ul |
| Donor DNA | 1ul | 60 ng/ul | 15 ng/ul (20-30ng/ul) |
| Phenol Red | 0.5ul | | |

2) Screening

The key to creating knock-in lines lies in simplifying the screening process for fish. If the desired fluorescent expression is not observed in a significant proportion of embryos injected with sgRNA + donor DNA + cas9, it may be necessary to revise your scheme. A distinct sgRNA site and/or adding a minimal promoter may be required. So, you require to prepare new donor DNA. Although this demands additional effort and time, it can lead to successfully create a line as injecting the same components may not yield to generate a line.

While screening, visual inspection holds greater importance. Selecting larvae with positive fluorescence is much easier than genotyping a large number of fish.

To increase the rate of selecting founders, double screening approach would help. First screening is to sort fluorescence-positive larvae during 0-5 dpf. Second screening is to select fluorescence -positive adult at 2-3 mpf.

In the case of injecting non-fluorescent DNA, such as CreER, it is recommended to include a selection marker in the donor plasmid. A selection marker can be franked by FRT, such as FRT-a-cry:venus-pA-FRT, allowing to remove it by injecting flp mRNA.

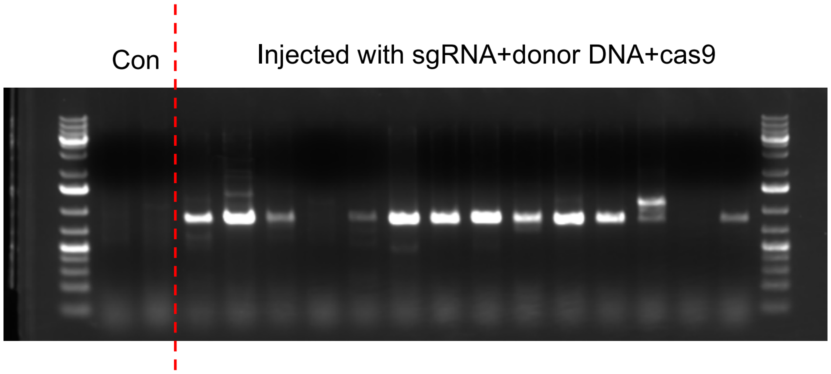


Figure: example of genotyping

Con: Control wild-type embryos
